# MAIT cells activate dendritic cells to promote T follicular helper cell differentiation and humoral immunity

**DOI:** 10.1101/2022.03.31.486638

**Authors:** Theresa E. Pankhurst, Kaitlin H. Buick, Joshua L. Lange, Andrew J. Marshall, Kaileen R. Button, Olga R. Palmer, Kathryn J. Farrand, Isabelle F. N. Stewart, Thomas Bird, Ngarangi C. Mason, Benjamin J. Compton, Davide Comoletti, Mariolina Salio, Vincenzo Cerundolo, Gavin F. Painter, Ian F. Hermans, Lisa M. Connor

**Affiliations:** School of Biological Sciences, Victoria University of Wellington; Wellington, New Zealand; Malaghan Institute of Medical Research; Wellington, New Zealand; Ferrier Research Institute, Victoria University of Wellington; Wellington, New Zealand; Medical Research Council Human Immunology Unit, Weatherall Institute of Molecular Medicine, University of Oxford; Oxford, United Kingdom

## Abstract

Protective immune responses against respiratory pathogens, including influenza virus are initiated by the mucosal immune system. However, most licensed vaccines are administered parenterally and are largely ineffective at inducing mucosal immunity. The development of safe and effective mucosal vaccines has largely been hampered by the lack of a suitable mucosal adjuvant. In this study we explore a novel class of adjuvant that harness mucosal-associated invariant T (MAIT) cells. We show evidence that intranasal immunisation of MAIT cell agonists co-administered with protein, including haemagglutinin from influenza A virus induced potent humoral immunity and immunoglobulin (Ig)A production, which protected mice against infection. MAIT cell adjuvant activity was mediated by CD40L-dependent activation of dendritic cells and subsequent priming of CD4^+^ T follicular helper cells. In summary, we show that MAIT cells are promising vaccine targets that can be utilised as cellular adjuvants in mucosal vaccines.

## Introduction

Most infectious agents invade the body through the mucosa, yet the anatomic compartmentalization of mucosal immunity is not incorporated into the design of most licensed vaccines, which are typically administered by parenteral routes - commonly by intramuscular or subcutaneous injection. While parenteral vaccination can be efficacious, notably where the infected tissues are permeable to transudation by serum antibodies, the mucosal tissues tend to be better served by immune responses provoked locally^1^. Mucosal vaccination, such as via oral and nasal routes, may therefore be more likely to elicit antigen-specific immune responses that home to mucosal tissues to neutralise or eliminate pathogens at the site of infection^1^. Such responses will include T cells and B cells that respond initially to vaccine antigens that drain to mucosa-associated lymphoid tissues; these cells then transit via the lymph and circulation to specifically seed mucosal sites as effector and memory cells. Importantly, mucosal delivery is more likely than parenteral administration to lead to mucosal IgA, a key immune factor responsible for preventing pathogen invasion of the host to cause infection^2^.

Currently the only approved vaccines delivered by the intranasal route (as nasal sprays) are the live attenuated influenza vaccines FluMist^3^ and Nasovac-S^4^. However, due to inconsistent protective efficacy, perhaps reflecting unwanted biasing of the immune response to previously encountered influenza strains, and because of safety concerns regarding the use of live viral strains, new subunit vaccines with refined specificity are needed^5^. As subunit vaccines typically require immune adjuvants to achieve efficacious responses, more work is required to develop safe and effective adjuvants that can be used by the nasal route.

Mucosal-associated invariant T (MAIT) cells are unconventional T cells with innate-like effector functions that are abundant in mucosal tissues^6,7^. While they have been shown to have anti-microbial activity, specifically by recognising non-protein agonists derived from vitamin B2 riboflavin synthesis pathways in some microbial species^8–12^, they also have broader immunomodulatory activities that could potentially be exploited in vaccination strategies. An attractive feature in this regard is the limited diversity of the MAIT T cell receptor (TCR), which has been highly conserved throughout mammalian evolution, and the lack of polymorphism of major histocompatibility class (MHC) 1-related molecule (MR1), the non-classical MHC class I molecule on which agonists are presented; MAIT cell activation can therefore be achieved in all individuals with the same agonist compounds. Agonists that have been identified include 5-(2-oxoethylideneamino)-6-D-ribitylaminouracil (5-OE-RU) and 5-(2-oxopropylideneamino)-6-D-ribitylaminouracil (5-OP-RU), which are compounds formed naturally through non-enzymatic condensation of the vitamin B pathway derivative 5-amino-6-D-ribitylaminouracil (5-A-RU) with glycolysis by-products glyoxal and methylglyoxal (MG) respectively^9^. In animal models, intranasal administration of MAIT cell agonists combined with toll-like receptor (TLR) agonists has shown to activate MAIT cells in the lung^13–14^. Importantly, in vitro studies with human cells have shown that activation of MAIT cells with 5-A-RU/MG can lead to enhanced dendritic cell (DC) function^15^. As activation of DCs is necessary for the development and differentiation of effector T cell responses, including the T follicular helper (T_FH_) cells required for B cell activation, formation of germinal centre (GC) reactions and production of high affinity antibodies^16^, it is possible that agonists such as 5-OP-RU could act as a mucosal adjuvants when delivered with antigen.

In this study we evaluated the capacity of 5-A-RU/MG to enhance immunogenicity of protein antigens when co-administered by intranasal instillation. We show that activation of MAIT cells by this route provides an adjuvant effect, promoting CD40L-dependent activation of lung-associated DCs, expansion of T_FH_ cells and production of antigen-specific mucosal IgA. When combined with a protein antigen from influenza A virus (IAV), complete protection from virus challenge was observed, highlighting the potential to utilise MAIT cells as “cellular adjuvants” in vaccine design.

## Results

### Intranasal co-administration of a MAIT cell agonist with an antigen enhances specific antibody response

Studies conducted in vitro have shown that MAIT cells are able to induce the production of antibodies by B cells^17^, however, the extent of their ability to promote such responses in vivo has not yet been investigated. To determine whether the activation of MAIT cells could promote humoral responses towards co-delivered antigen, we established an immunisation regimen to measure antigen-specific B cells and antibody responses (Fig. 1a). Intranasal (i.n.) administration of 5 nmol EndoGrade® ovalbumin (OVA) protein alone, or in combination with 75 nmol of freshly prepared 5-OP-RU (a 1:10 ratio of 5-A-RU and MG mixed immediately prior to use, hereafter referred to as “5-A-RU/MG”) was given to mice three times at two-week intervals. One week following the third dose, the proportions of OVA-specific B cells were determined in the lung-draining mediastinal lymph nodes (mLN) by flow cytometry using biotinylated OVA protein. The levels of OVA-specific antibodies in the serum, and mucosal antibodies in the bronchoalveolar lavage (BAL) were also assessed by enzyme-linked immunosorbent assay (ELISA). As seen in Fig. 1b-c, the total frequency and number of GC B cells (B220^+^ GL7^+^ IgD^−^) was significantly greater in wild type (WT) C57BL/6 mice treated with 5-A-RU/MG + OVA compared to mice treated with OVA alone, and the proportion of OVA-specific (B220^+^ GL7^+^ OVA-biotin^+^) B cells was significantly enhanced in this group. In contrast, there was no expansion of OVA-specific B cells in MR1^−/−^ mice (that lack MAIT cells) treated with 5-A-RU/MG + OVA, confirming the required role of MAIT cells in this response (Fig. 1c). The detection of OVA-specific B cells by flow cytometry correlated with the presence of circulating OVA-specific serum IgG antibodies, which was also dependent on the presence of MR1 (Fig. 1c). The antibody responses induced by MAIT cell agonist treatment were dominated by IgG1, whereas minimal levels of IgG2a/c, IgG2b or IgG3 were detected. Serum IgE was not generated after intranasal 5-A-RU/MG + OVA, whereas serum taken from mice infected with *Schistosoma mansoni* had high IgE levels, as expected for this positive control (Fig. 1e). Given that secretory IgA (sIgA) present in the airway lumen provides neutralising activity in mucosal sites^18^, it was notable that levels of OVA-specific sIgA were significantly enhanced in mice treated with 5-A-RU/MG + OVA (Fig. 1f). Collectively, these data suggest that MR1-restricted MAIT cells have the capacity to promote mucosa-associated antigen-specific B cells, serum IgG and mucosal sIgA antibodies; agonists for MAIT cells can therefore be considered as mucosal vaccine adjuvants.

**Fig. 1.**
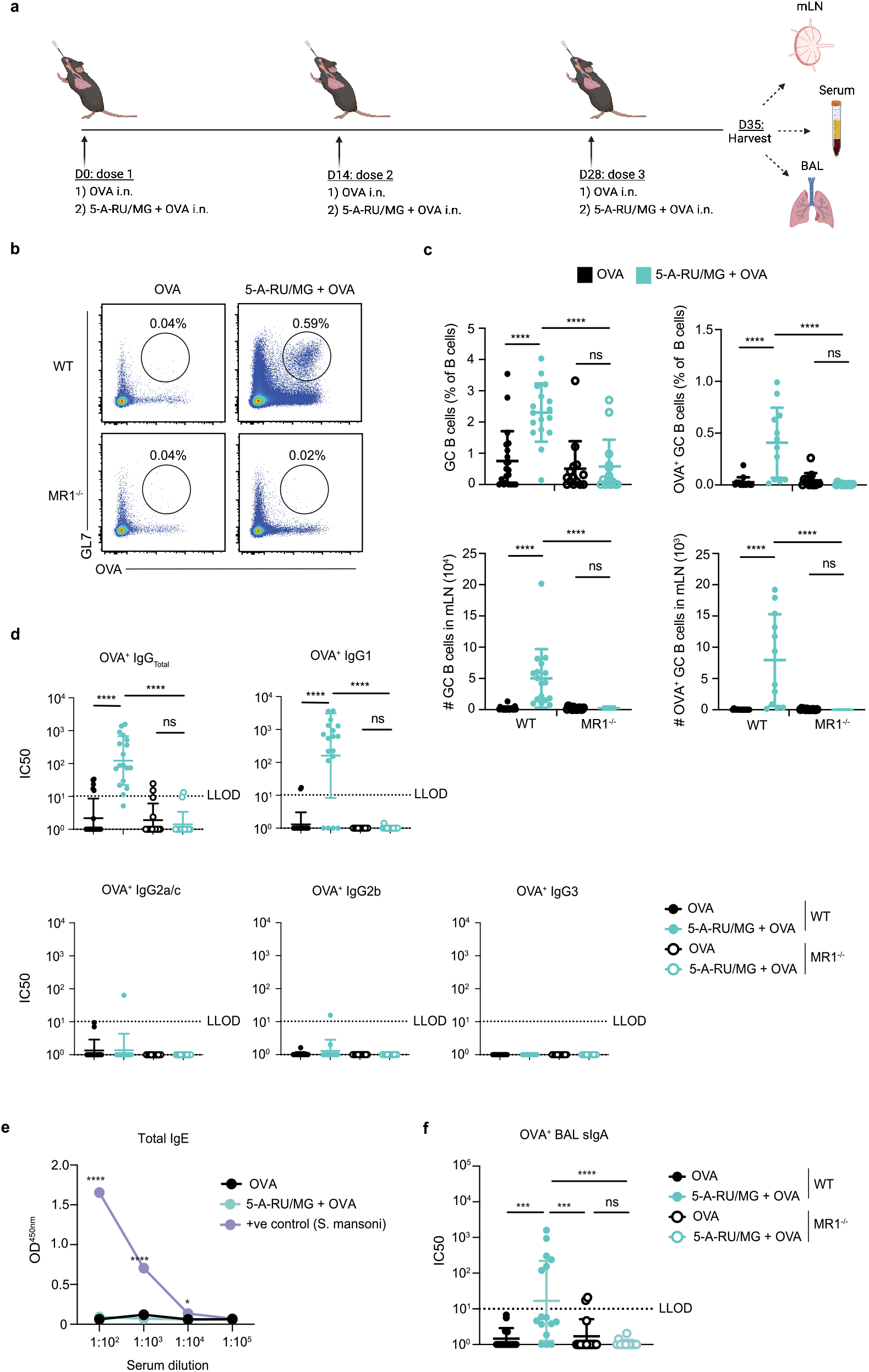
Intranasal co-administration of a MAIT cell agonist with an antigen enhances specific antibody response. (**a**) WT or MR1^−/−^ mice were treated i.n. with either 5 nmol OVA alone or in combination with 75 nmol 5-A-RU and 750 nmol MG, three times spaced two weeks apart. One week after the final booster dose, mLN, serum and BAL were harvested for analysis. Graphic created at Biorender.com. (**b**) Representative flow cytometry plots of mLN OVA-specific GC B cells (TCRβ^−^ B220^+^ GL7^+^ OVA-biotin-SAV^+^). Full gating strategy available in Supplementary Figure 1. **(c)** Frequency and total number of GC B cells and OVA-specific GC B cells in the mLN. (**d**) Half maximal inhibitory concentration (IC50) Log10 values from serum OVA-specific IgG ELISA. (**e**) Optical density (OD; 450nm) from total IgE serum ELISA **(f)** IC50 from OVA-specific BAL sIgA ELISA. Lower limit of detection (LLOD) was set at an IC50 of 10, IC50 values below zero were marked as an IC50 of one. Data represent a combination of *n*=3 (**b-d**) and *n*=2 (**e-f**) individual experiments, with *n*=3-10 mice per group, per experiment. Graphs depicted as mean ± SD. One-way (**c, d, f**) or Two-way (**e**) ANOVA with Tukey’s multiple comparisons test was performed with ns>0.05, \**P*≤0.05, \*\*\**P*≤0.001, \*\*\*\**P*≤0.0001.

### Intranasal co-administration of a MAIT cell agonist with a viral antigen induces protective response

To determine whether the antigen-specific antibodies generated by protein adjuvanted with 5-A-RU/MG have the potential to provide a protective role, humoral immunity against IAV haemagglutinin (HA; A/X-31) protein was assessed. HA is a glycoprotein expressed by the virion membrane of the influenza virus, which permits viral entry into host cells^19^. Thus, antibodies directed against HA protein can block viral infection and reduce disease, providing a functional readout of the immune response. To examine whether MAIT cells can help stimulate HA-specific antibodies, C57BL/6 (WT) mice were immunised i.n. with HA protein alone or in combination with 5-A-RU/MG as outlined in Fig. 2a. As a positive control for this response, an additional group of mice were treated subcutaneously (s.c.) with HA protein and the licensed adjuvant AddaVax. One week after the final boost, significantly enhanced levels of HA-specific IgG were observed in the serum of mice immunised with HA adjuvanted with 5-A-RU/MG or AddaVax when compared to HA alone (Fig. 2b). The IgG1 isotype dominated the response in both adjuvanted groups, with lower titres of HA-specific IgG2a/c and IgG2b (Fig. 2b). Titres of sIgA in the BAL fluid were enhanced with 5-A-RU/MG treatment compared to unadjuvanted groups (Fig. 2c) and were similar to levels of sIgA produced by mice previously infected with the X31 strain of IAV, a known inducer of mucosal sIgA^20^. Interestingly, s.c. delivered AddaVax + HA treatment generated moderate levels of sIgA, albeit almost a log-fold less than i.n. 5-A-RU/MG + HA treatment.

**Fig. 2.**
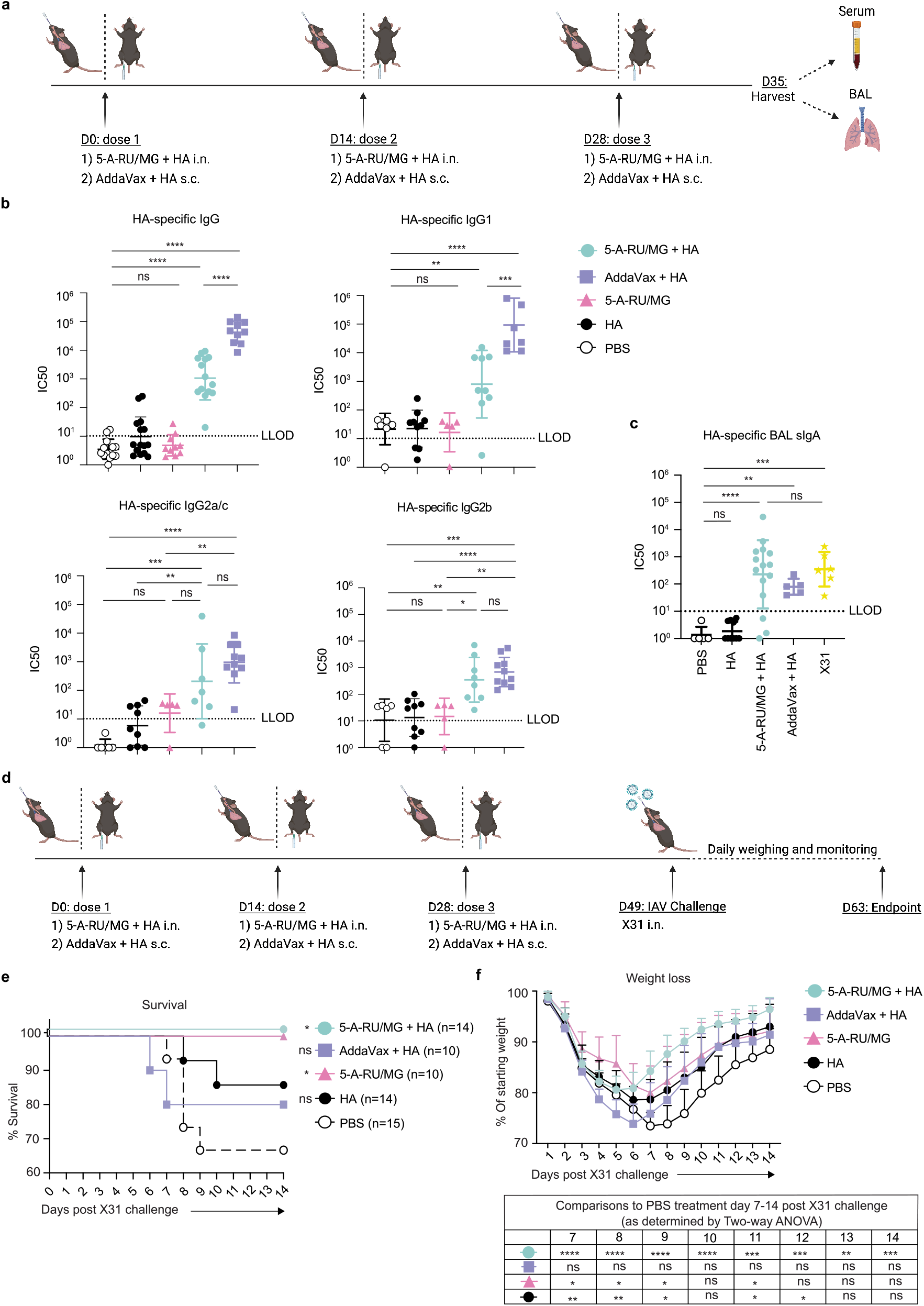
Intranasal co-administration of a MAIT cell agonist with a viral antigen induces protective response. **(a)** Schematic of experimental, created at Biorender.com.. WT mice were treated i.n. with 10 μg of HA protein alone or admixed with 75 nmol 5-A-RU and 750 nmol MG. AddaVax + 10 μg HA was delivered s.c. at 1:1 v/v. (**b**) IC50 Log10 values from HA-specific serum IgG ELISA. LLOD was set at an IC50 of 10, IC50 values below zero were marked as an IC50 of one (**c**) IC50 Log10 values from BAL HA-specific sIgA ELISA. (**d**) Schematic of experimental design, created at Biorender.com. All mouse groups were challenged with 2 HAU of X31 IAV. (**e**) Survival and (**f**) Weight loss. Data represent a combination of *n*=3 (**b, e, f**), or *n*=2 (**c**) individual experiments, with *n*=4-10 mice per group, per experiment. Graphs depicted as mean ± SD. Statistical significance was determined by One-way ANOVA (**b, c**), Kaplan-Meier survival curve comparison (**e**), or Two-way ANOVA (**f**) with Tukey’s multiple comparisons test (**b, c, f**) and log-rank (Mantel-cox) test (**e**). Comparisons displayed in (**e**) compare each treatment group to PBS treatment. Comparisons displayed in the table in (**f**) compare each treatment group to PBS treatment at day 7-14 post X31 challenge. ns>0.05 \**P*≤0.05, \*\**P*≤0.01, \*\*\**P*≤0.001, \*\*\*\**P*≤0.0001.

Immunised mice were then assessed for vaccine-induced protection against live influenza virus infection. Three weeks after the last boost, mice were challenged i.n. with 2 haemagglutinating units (HAU) of X31 IAV and monitored for weight loss (Fig. 2d). Animals with weight loss exceeding 30% of their initial weight were culled. Because activation of MAIT cells can stimulate a direct antiviral response^21–23^, a group of mice that had received 5-A-RU/MG without HA protein (according to the same administration regimen) was included for comparison. Indeed, treatment with the MAIT cell agonist alone proved to be sufficient to confer protection, with all animals surviving virus challenge (Fig. 2e). However, it was notable that mice treated with HA together with the MAIT cell agonist gained weight at a faster rate (Fig. 2f), likely reflecting the induced HA-specific adaptive response; again, all animals survived infection (Fig. 2e). Surprisingly, mice immunised s.c. with HA protein and AddaVax lost weight more rapidly than the other treatment groups and almost 20% of animals did not survive the infection (Fig. 2e-f), suggesting that the mucosal route of immunisation can provide enhanced protection against live influenza infection.

### Intranasal administration of agonist evokes MAIT cell activation and proliferation in the lung

To gain a better understanding of the role for MAIT cells during the intranasal immunisation regimen, we examined the response of lung-associated MAIT cells to increasing concentrations of 5-A-RU/MG. Mice were intranasally administered either 5, 20, 75, 140 or 180 nmol 5-A-RU, in each case mixed with 10-fold MG directly before use to form 5-OP-RU. Lungs and mLNs were isolated 24 hours later to assess MAIT cells by flow cytometry with fluorescent MR1/5-OP-RU tetramers and antibodies against CD69. All doses induced upregulated expression of CD69, indicative of MAIT cell activation, with the dose response levelling off at 75 nmol in both the lung and mLN (Fig. 3a-b).

**Fig. 3.**
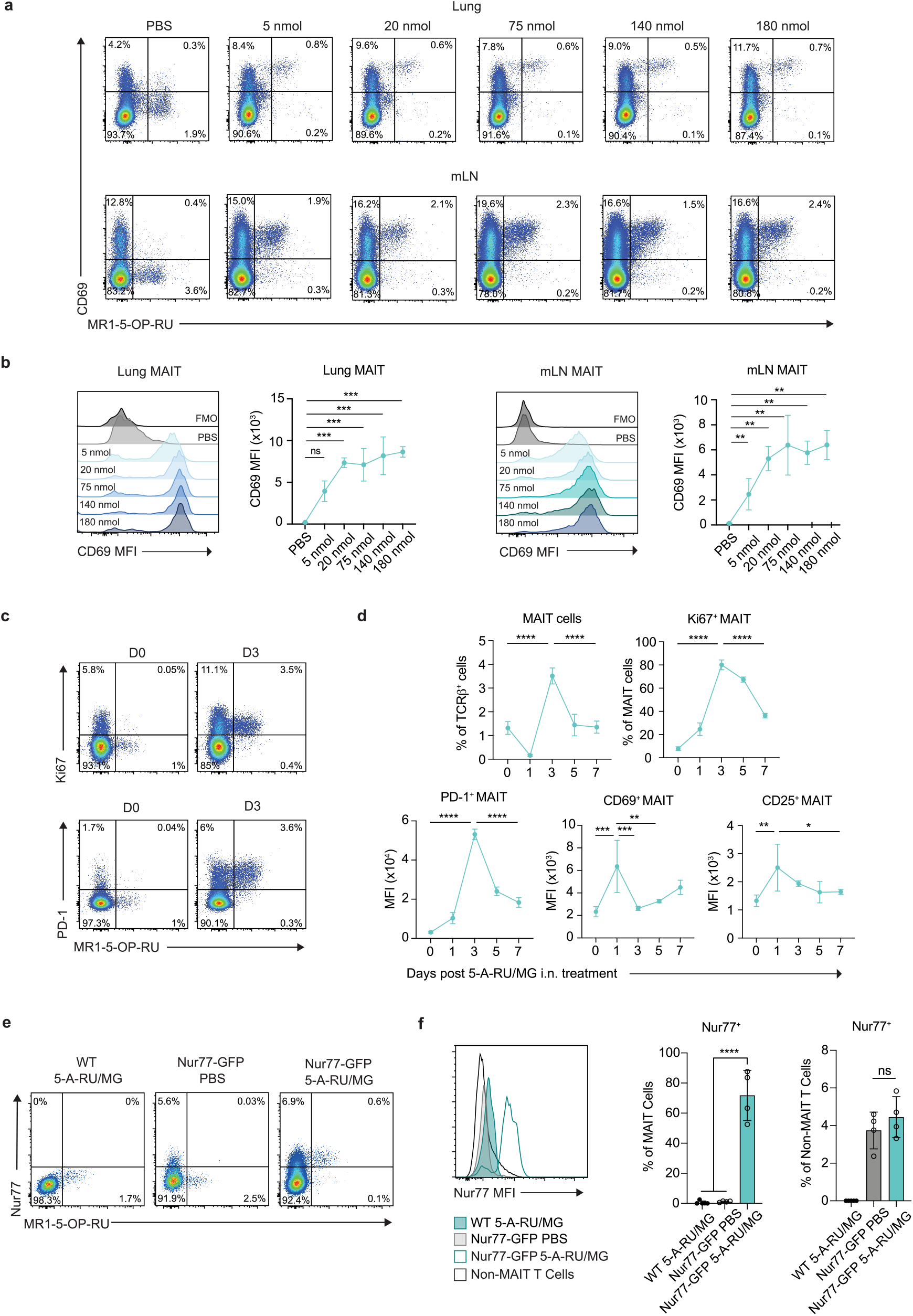
Intranasal administration of 5-A-RU/MG evokes MAIT cell proliferation and accumulation in the lung. (**a**) WT mice were treated i.n. with PBS or increasing doses of 5-A-RU with 10x the corresponding amount of MG. Lung and mLNs were collected 24 hours later for flow cytometry. Representative flow plots of CD69^+^ MAIT cells (TCRβ^+^B220^−^CD64^−^MR1-5-OP-RU tetramer^+^). (**b**) CD69 expression on MAIT cells. (**c**) WT mice were treated with 75 nmol 5-A-RU and 750 nmol MG i.n. and lungs were collected at indicated timepoints. Naïve mice were used as baseline day 0. Representative flow plots of Ki67^+^ and PD-1^+^ lung MAIT cells. (**d**) Frequency of total and Ki67^+^ MAIT cells in lung. MFI of PD-1, CD69 and CD25 on lung MAIT cells over time. (**e**) WT and Nur77-GFP mice were treated with PBS or 5-A-RU/MG i.n. and mLNs were collected 24 hours later. Representative flow plots of Nur77^+^ MAIT cells (TCRβ^+^B220^−^CD64^−^MR1-5-OP-RU tetramer^+^Nur77^+^). (**f**) MAIT cell (TCRβ^+^MR1-5-OP-RU tetramer^+^) and non-MAIT T cell (TCRβ^+^MR1-5-OP-RU tetramer^−^) Nur77 expression and frequency. Data is representative of *n*=2 individual experiments, with *n*=2-5 mice per group. Graphs depicted as mean ± SD. One-way ANOVA with Tukey’s multiple comparisons test was performed with ns>0.05, \**P*≤0.05, \*\**P*≤0.01, \*\*\**P*≤0.001, \*\*\*\**P*≤0.0001.

To examine kinetics of the response, mice were treated with 75 nmol 5-A-RU/MG (a dose that led to maximal CD69 upregulation) and MAIT cell phenotype was assessed in the lung tissue over a period of seven days (Fig. 3c-d). After 24 hours, a significant reduction in MAIT cell frequency was observed, consistent with previous reports showing downregulation of the MAIT cell TCR upon initial stimulation^14^. Despite this reduced capacity to detect MAIT cells, the cells that could be identified showed clear evidence of activation, with significantly enhanced expression of CD69 and CD25, with levels waning over the following days. Expression of programmed cell death protein 1 (PD-1) also increased but peaked later at day three (Fig. 3d). The number of MAIT cells detected in the lung peaked at day three and reduced to baseline levels by day five, with peak accumulation coinciding with evidence of proliferation as indicated by Ki67 expression (Fig. 3d). To confirm that MAIT cells were being stimulated through their invariant TCR via MR1 presentation of agonists, we employed Nur77^GFP^ mice which express green fluorescent protein (GFP) in response to TCR stimulation in a dose-dependent manner^24^. Induced Nur77 expression was detected on 70% of MAIT cells in the mLN at 24 hours following i.n. 5-A-RU/MG administration, representing a 60-fold change relative to PBS-treated controls (Fig. 3e-f). In contrast, there were minimal changes in Nur77 expression of non-MAIT T cells (Fig. 3f). Overall, these data show that i.n. 5-A-RU/MG can induce MAIT cell activation, proliferation, and accumulation in the lung and mLN, attributable (at least in part) to direct stimulation via the TCR.

### Intranasal co-administration of 5-A-RU/MG + OVA induces OVA-specific T_FH_ cell development

Given the essential role T_FH_ cells play in humoral immune responses, the induction of this population is an important process in vaccination. To investigate the T_FH_ response induced after i.n. 5-A-RU/MG treatment, expression levels of the T_FH_-associated molecules B-cell lymphoma 6 protein (Bcl6), PD-1, folate receptor 4 (FR4) and C-X-C motif chemokine receptor 5 (CXCR5) were assessed in the mLN. As MAIT cells share some similarities with natural killer T (NKT) cells, and the latter are capable of adopting a T_FH_-like phenotype and provide direct help to B cells^25–26^, analysis was conducted on both CD4^+^ T cell and MAIT cell populations.

As MAIT cell activation and accumulation occurred within the first three days following 5-A-RU/MG treatment, whereas conventional T cell responses typically require more time to achieve a measurable response^27^, the expression of T_FH_-associated molecules was assessed at early (day three) and later (day seven) timepoints. The transcription factor Bcl6, which is necessary for T_FH_ differentiation was only detected within the non-MAIT (negative for MR1-5-OP-RU tetramer) T cell population (Fig. 4a). Furthermore, expression of inducible co-stimulator (ICOS), CXCR5 and FR4 was not upregulated on MAIT cells at day three or seven after 5-A-RU/MG treatment (Fig. 4b). Among conventional CD4^+^ T cells, expanded populations of Bcl6^+^ PD-1^+^ cells were detected in the mLN at day seven (Fig. 4c), suggesting immunisation with 5-A-RU/MG promotes the induction of conventional CD4^+^ T_FH_ cells. In contrast, Bcl6^+^PD-1^+^ MAIT cells did not expand (Fig. 4c). To determine whether any of the conventional T_FH_ cells induced were OVA-specific, CD4^+^ T cells with specificity for (one or two) dominant epitopes were identified using a combination of two MHC class II tetramers, containing I-A(b)HAAHAEINEA and I-A(b)AAHAEINEA epitopes respectively (that were labelled with the same fluorophore), and then assessed for Bcl6 and PD-1 expression. In vaccinated WT mice, OVA-specific T cells represent on average 0.15% of all CD4^+^ T cells, and of these an average of 20% expressed a T_FH_ phenotype (Fig. 4d-e). The development of OVA-specific T_FH_ cells was significantly reduced in unadjuvanted WT mice and in MR1-deficient mice (Fig. 4d-e), suggesting that mucosal MAIT cell stimulation can drive conventional CD4^+^ T_FH_ differentiation against co-delivered proteins.

**Fig. 4.**
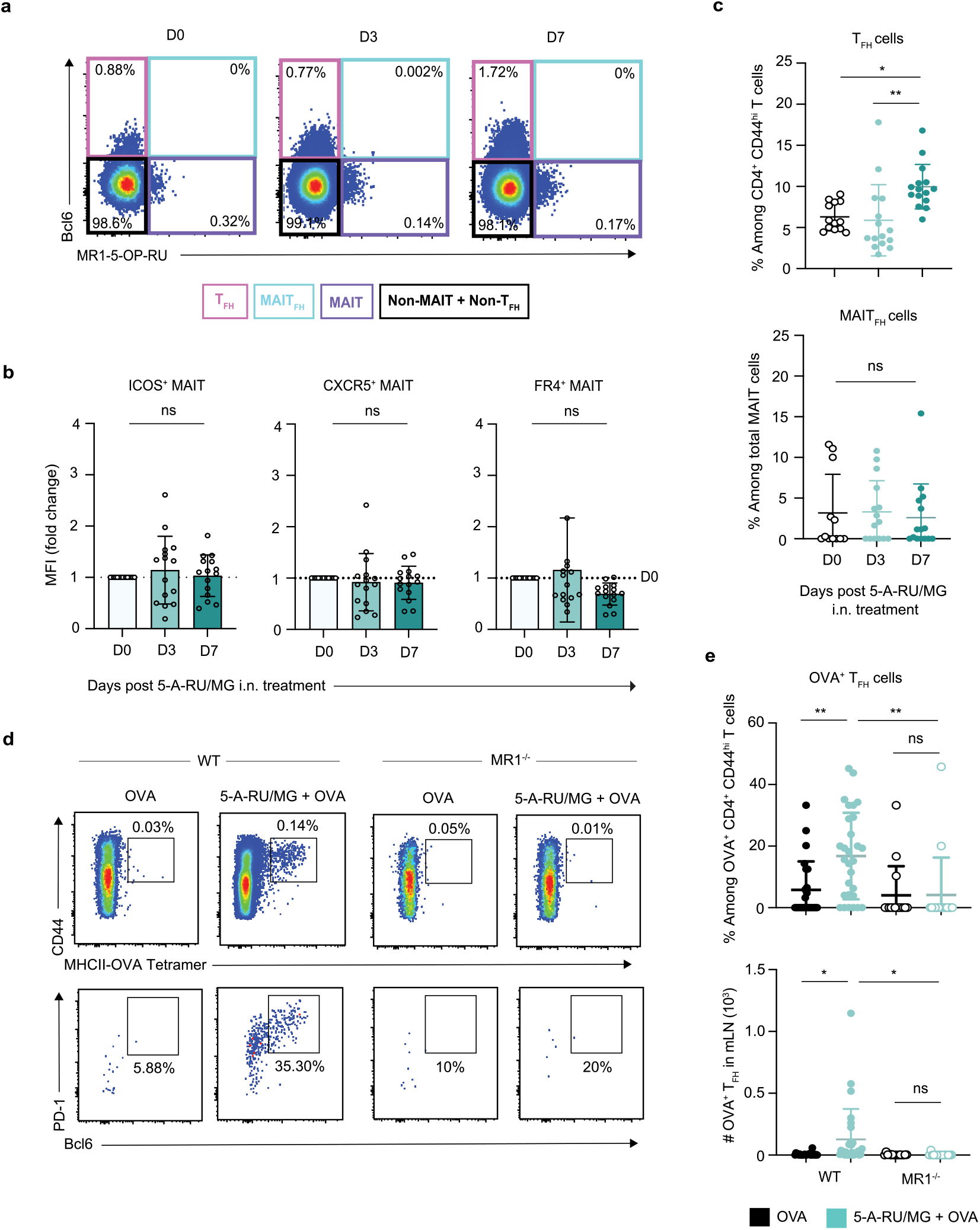
Intranasal co-administration of 5-A-RU/MG + OVA promotes generation of conventional OVA-specific TFH cells. WT mice were treated i.n. with a single dose of 75 nmol 5-A-RU and 750 nmol MG. Naïve (D0) and treated mLN were harvested three and seven days later for flow cytometry. Full gating strategy available in Supplementary Figure 2. (**a**) Representative flow cytometry plots of T cells (TCRβ^+^) expressing MR1-5-OP-RU and Bcl6. (**b**) MFI fold change compared to D0 mice of T_FH_ markers ICOS, CXCR5 and FR4 expressed by MAIT cells. (**c**) Frequency of conventional T_FH_ (B220^−^TCRβ^+^CD4^+^CD44^hi^Bcl6^+^PD1^+^) and MAIT_FH_ (B220^−^TCRβ^+^CD64^−^MR1-5-OP-RU^+^Bcl6^+^). (**d**) WT and MR1^−/−^ mice were treated i.n. with three doses of either OVA alone or in combination with 5-A-RU/MG, spaced two weeks apart. mLN were harvested five days after the final boost for analysis of TFH by flow cytometry. Representative flow plots of OVA-specific CD4^+^ T cells and OVA-specific TFH cells. (**e**) Frequency and total numbers of OVA-specific TFH in the mLN. Data represent a combination of *n*=3 (**a-c**), *n*=4 (**d-e**) individual experiments, with *n*=3-10 mice per group, per experiment. Graphs depicted as mean ± SD. One-way ANOVA with Tukey’s multiple comparisons test was performed with ns>0.05, \**P*≤0.05, \*\**P*≤0.01.

### Conventional CD4^+^ T_FH_ cells are essential for humoral immunity induced by 5-A-RU/MG + OVA

To confirm the role of CD4^+^ T_FH_ cells in the induction of humoral immunity by treatment with 5-A-RU/MG + OVA, we next assessed the adaptive immune response generated by i.n. immunisation in B6Aa0^−/−^ mice, which lack MHC class II molecules and do not produce conventional CD4^+^ T cells. While B6Aa0^−/−^ mice could raise antibody responses specific for T-independent hapten antigen 4-hydroxy-3-nitrophenyl (Supplementary Fig. 3), they failed to induce OVA-specific GC B cells (Fig. 5a-b) or OVA-specific serum IgG antibodies (Fig. 5c) in response to 5-A-RU/MG + OVA treatment. Importantly, MAIT cell activation remained intact in B6Aa0^−/−^ mice, displaying comparable frequency of MAIT cells expressing CD69 and PD-1 to C57BL/6 mice (Fig. 5d). Taken together, these data show a critical role for conventional T_FH_ cells in the antigen-specific humoral immune response generated by a MAIT cell-based adjuvant mucosal vaccine, that activated MAIT cells could not achieve alone.

**Fig. 5.**
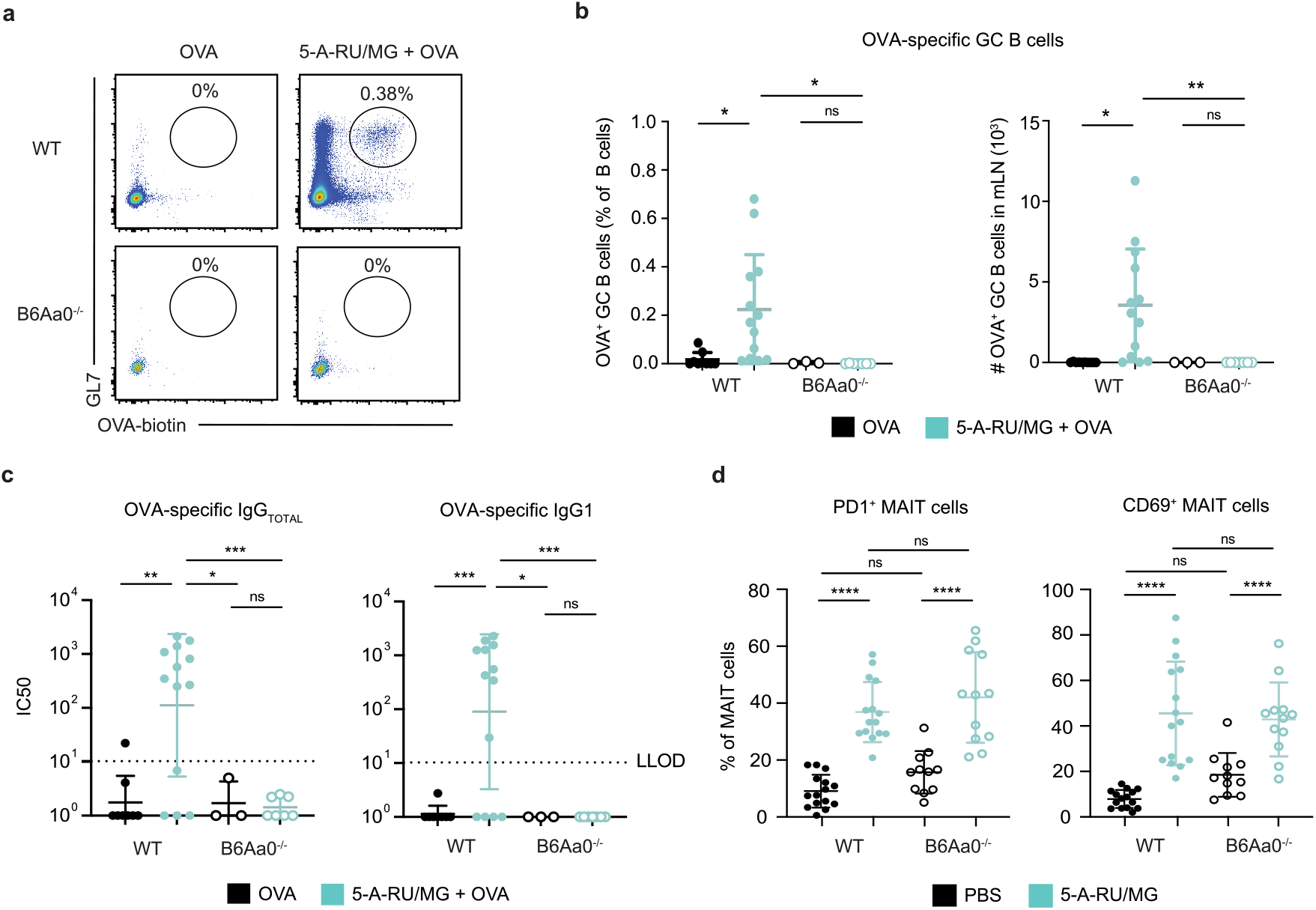
Conventional CD4 TFH cells are essential for humoral immunity induced by 5-A-RU/MG + OVA immunisation. WT or B6Aa0^−/−^ mice were treated i.n. with either 5 nmol OVA alone or in combination with 75 nmol 5-A-RU and 750 nmol MG, three times spaced two weeks apart. One week after the final booster dose, mLN and serum were harvested for analysis. (**a**) Representative flow cytometry plots of OVA-specific GC B cells (TCRβ^−^ B220^+^GL7^+^OVA-biotin-SAV^+^). (**b**) Frequencies and total numbers of OVA-specific GC B cells in the mLN. (**c**) IC50 Log10 values from OVA-specific serum IgG ELISA. LLOD was set at an IC50 of 10, IC50 values below zero were marked as an IC50 of one (**d**) WT and B6Aa0^−/−^ mice were treated i.n. with a single dose of PBS or 5-A-RU/MG. 24 hours later lungs were harvested for flow cytometry. Frequency of PD1 and CD69 expressing MAIT cells (B220^−^ TCRβ^+^CD64^−^MR1-5-OP-RU^+^). Data represent a combination of *n*=2 (**a-c**) **or** *n*=3 (**d**) individual experiments, with *n*=3-8 mice per group, per experiment, apart from OVA-only treated B6Aa0^−/−^ mice in B-C, that only had 3 mice total. One-way ANOVA with Tukey’s multiple comparisons test was performed with ns>0.05, \**P*≤0.05, \*\**P*≤0.01, \*\*\**P*≤0.001, \*\*\*\**P*≤0.0001.

### Intranasal administration of 5-A-RU/MG activates migratory DCs

DCs are the vital link between the innate and adaptive immune response and are essential for priming CD4^+^ T_FH_ cells^28–30^. DCs possess the specialised antigen presenting machinery necessary for MAIT cell activation, and in turn, studies have shown that activated MAIT cells can provide reciprocal stimulatory signals that mature the DC in vitro^15^. The accumulation of the different conventional DC (cDC) subsets was therefore assessed one day following i.n. 5-A-RU/MG administration. In the mLN, resident DCs can be identified as MHCII^Int^CD11c^+^ cells, while migratory DCs can be identified as MHCII^hi^CD11c^+^ cells, the latter representing recent migrants from the lung tissue. Both of these cDC subsets can be further divided into the functionally distinct populations cDC1 (CD11c^+^signal regulatory protein α (SIRPα)^−^) and cDC2 (CD11c^+^CD11b^+^SIRPα^+^) which are responsible for the differentiation of distinct T cell populations^31–33^. Using the gating strategy in Supplementary Fig. 4 (and summarized in Fig. 6a), we assessed the impact of i.n. 5-A-RU/MG administration on cell number and phenotype of these different DC populations. No significant changes in cell number or activation status were observed for the resident populations (Fig. 6b and Supplementary Fig. 5). In contrast, the frequency of migratory DC was increased (with a non-significant upward trend in cell number) (Fig. 6c), and activation markers CD80, CD86 and programmed cell death ligand 1 (PDL-1) were also upregulated (Fig. 6d), observed on both cDC1 and cDC2 subsets (Fig. 6e). Importantly, no change in activation status of these DCs was seen when 5-A-RU/MG was administered to MR1^−/−^ mice, but was observed when administered to B6Aa0^−/−^ mice (Fig. 6f), highlighting the important MAIT cell interaction required.

**Fig. 6.**
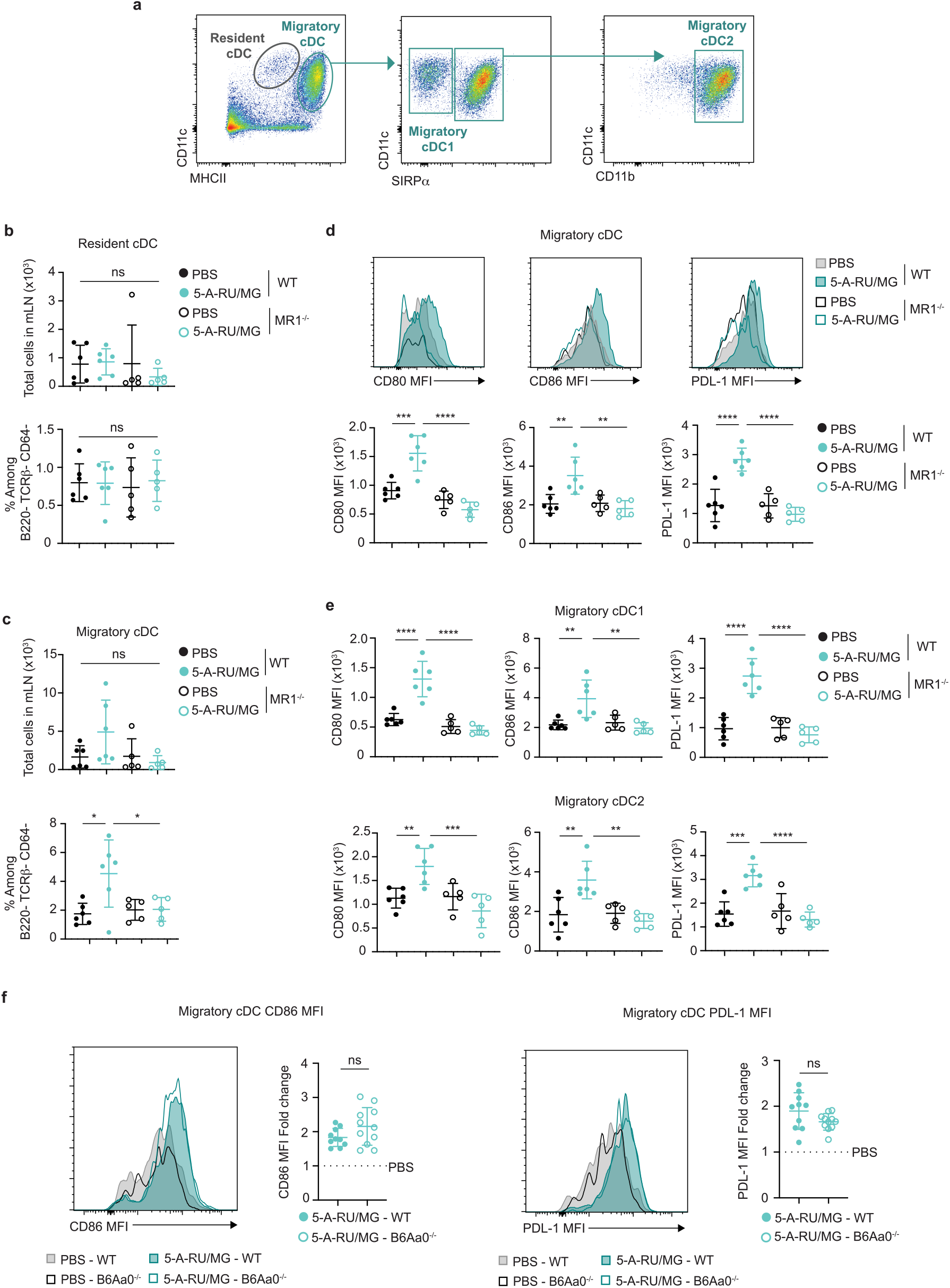
Intranasal administration of 5-A-RU/MG activates migratory dendritic cells. WT or MR1^−/−^ mice were administered i.n. with PBS or 75 nmol 5-A-RU and 750 nmol MG. 24 hours later mLN were harvested for flow cytometry. (**a**) Gating strategy to identify cDC subsets. (**b**) Frequency and total numbers of resident cDCs. **(c)** Frequency and total numbers of migratory cDCs. (**d**) Representative histograms and MFI of activation markers expressed by migratory cDCs. (**e**) MFI of activation markers expressed by migratory cDC1 and migratory cDC2 subsets. (**f**) WT or B6Aa0^−/−^ mice were administered and harvest as above. Representative histograms and fold change of CD86 and PDL-1 MFI expressed by migratory cDC. Fold change is relative to the average of PBS treated mice for each mouse strain. Data is a representative (a**-e**) or a combination (**f**) of *n*=2 independent experiments, with *n*=5-6 mice per group, per experiment. Graphs depicted as mean ± SD. Statistical significance was determined by One-way ANOVA with Tukey’s multiple comparisons test (**b-e**) or unpaired t-test (**f**), ns>0.05, \**P*≤0.05, \*\**P*≤0.01, \*\*\**P*≤0.001, \*\*\*\**P*≤0.0001.

### MAIT cell activation of cDCs is dependent on CD40-CD40L interaction

Signaling via CD40 is a well-known pathway of DC activation, resulting in increased expression of co-stimulatory molecules and release of proinflammatory cytokines, ultimately facilitating initiation of T cell immunity to acquired antigens^34^. Moreover, CD40-CD40L ligation was shown to be essential for DC activation in vitro following incubation with CD40L-expressing human MAIT cells^15^. It is therefore notable that we detected significant upregulation of CD40L on MAIT cells from the lung within three hours after i.n. 5-A-RU/MG treatment compared to MAIT cells from PBS treated mice (Fig. 7a). To determine whether CD40 ligation was necessary for MAIT cell-dependent DC activation, we used a monoclonal anti-CD40L blocking antibody (αCD40L mAb) to prevent CD40:CD40L signaling during 5-A-RU/MG treatment, and then assessed levels of MAIT cell activation and phenotype of migratory cDC (Fig. 7b). Blocking CD40 signaling did not affect MAIT cell activation, as indicated by CD69 upregulation (Fig. 7c). In contrast, upregulation of the co-stimulatory molecule CD86 on migratory cDC1 and cDC2 subsets was inhibited when the blocking antibody was used (Fig. 7c), suggesting that CD40L signaling is essential for MAIT cells to activate migratory cDCs.

**Fig. 7.**
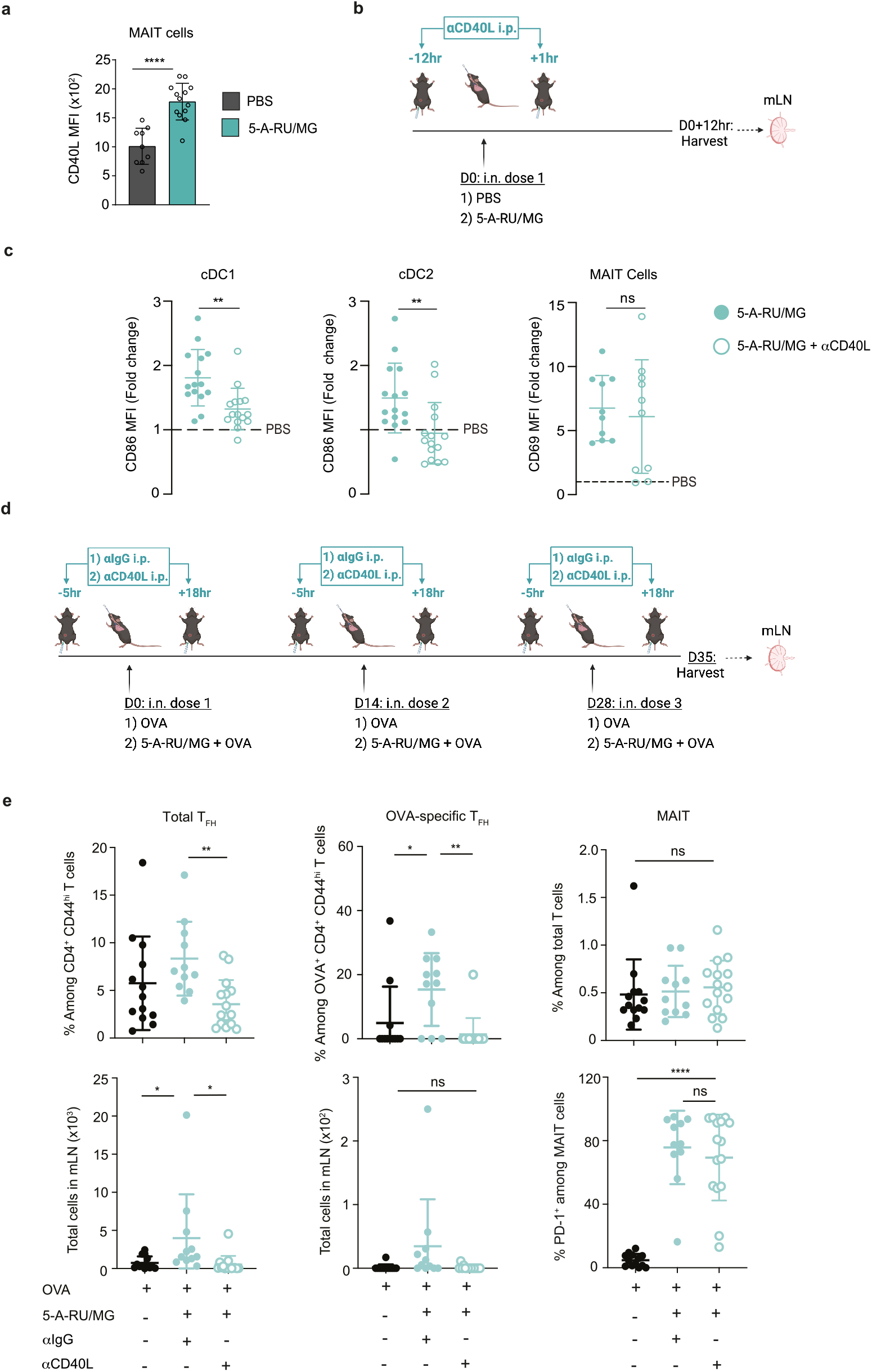
CD40L signaling is required for migratory cDC maturation and downstream induction of TFH after intranasal 5-A-RU/MG administration. (**a**) WT mice were treated i.n. with either PBS or 75 nmol 5-A-RU and 750 nmol MG. Three hours later, lungs were harvested for flow cytometry. MFI of CD40L expression by MAIT cells. (**b**) Schematic of experimental design, created at Biorender.com. 12 hours prior and 1 hour after i.n. treatment, mice were given 500 μg aCD40L i.p. (**c**) Fold change MFI of CD86 on migratory cDC1 (B220^−^TCRβ^−^CD11c^+^MHCII^hi^SIRPα^−^) and cDC2 (B220^−^TCRβ^−^ CD11c^+^MHCII^hi^SIRPα^+^CD11b^+^). Fold change of CD69 MFI on MAIT cells (B220^−^ TCRβ^+^CD64^−^MR1-5-OP-RU^+^). Fold change is relative to the average of PBS treated mice. (**d**) Schematic of experimental design, created at Biorender.com. WT mice were treated i.n. with either 5 nmol OVA alone or in combination with 5-A-RU/MG, three times spaced two weeks apart. Five hours prior and 18 hours after i.n. treatment, mice were given 500 μg aCD40L i.p. (**e**) Frequency and total number of mLN T_FH_ cells, OVA-specific T_FH_ cells, MAIT cells. Graphs depicted as mean ± SD. Data represent a combination of *n*=2 (**a**) or *n*=3 (**c, e**) individual experiments, with *n*=3-8 mice per group, per experiment. Unpaired t-test (**a, c**), or one-way ANOVA with Tukey’s multiple comparisons test (**e**) were performed with ns>0.05, \**P*≤0.05, \*\**P*≤0.01, \*\*\*\**P*≤0.0001

To establish whether CD40L-dependent activation of migratory cDC by MAIT cells was important for T_FH_ priming, CD4^+^ T cell responses were assessed following short-term αCD40L mAb blockade during treatment with 5-A-RU/MG + OVA. The antibody was administered five hours before and 18 hours after each of the three i.n. doses administered; a control group received an αIgG isotype control as outlined in Fig. 7d. Blocking CD40 signaling abolished the adjuvant effect of 5-A-RU/MG on T_FH_ development, with antigen-specific T_FH_ frequency and number reduced to levels seen with injection of OVA alone. Agonists for MAIT cells therefore support humoral responses to mucosally administered antigens by activating migratory cDC in a CD40-dependent fashion, leading to priming of conventional T_FH_ cells.

## Discussion

The results presented here show that activation of lung-associated MAIT cells by intranasal administration of MR1-binding agonists enhances humoral responses to co-administered antigens. This adjuvant activity involves CD40 signaling between activated MAIT cell and lung-associated migratory DCs promoting CD4^+^ T_FH_ cell priming and the development of protein-specific antibody responses. Notably, i.n. administration of 5-OP-RU (formed by admixing 5-A-RU with MG), and HA protein from IAV protected mice from lethal challenge, outperforming parenteral vaccination with a benchmark oil-in-water adjuvant.

DC licensing was a term first used to describe the action of CD4^+^ T cells to enhance DC activation, which in turn primed potent, long-lived CD8^+^ T cell responses^35^. Initially, provision of CD40L to activate DCs was thought to be a function primarily reserved for CD4^+^ T cells^36–38^. However, there is a growing body of evidence to suggest that innate-like T cell populations including NKT cells^39,40^, γ/δ T cells^41^ and MAIT cells^15^ can also perform this role. Here we report evidence for MAIT cell-dependent, CD40-mediated, DC activation in vivo. Like activated CD4^+^ T cells, we found that MAIT cells also express high levels of CD40L early after activation with 5-A-RU/MG treatment. Importantly, blocking CD40/CD40L interactions resulted in reduced expression of co-stimulatory molecules on DC, and complete ablation of T_FH_ CD4^+^ differentiation, suggesting a non-redundant role for CD40 signaling in the cellular adjuvant activity of MAIT cells. In DCs, signaling induced by pattern recognition (or via some cytokine receptors) is required to activate canonical nuclear factor kappa-light-chain-enhancer of activated B cells (NF-κB), which triggers alternative activation pathways, such as the interferon (IFN) response^42^. In the absence of these activating signals, the CD40 signaling pathway initiates a separate signaling cascade to activate NF-κB2^43^, which is typical of the non-canonical NF-κB signaling pathway^44^. The induction of MAIT cell-mediated CD40 signaling in the absence of pattern recognition signals, may explain the non-redundant role for CD40 signaling observed in this study. We aimed to block CD40 signaling activity during early MAIT cell-DC interactions by restricting αCD40L mAb administration to the priming phase. Nonetheless, the involvement of CD40-CD40L signaling on processes downstream of DC activation cannot be ruled out, such as CD40-CD40L interaction between DCs and naïve CD4^+^ T cells, which has been linked to T_FH_ differentiation^29^ and direct T cell help to B cells^45^.

The subset of DCs presenting antigen to naïve CD4^+^ T cells has been implicated in the instruction of Th differentiation including T_FH_ CD4^+^ differentiation^46^. While both migratory cDC and resident cDC have been shown to be capable of priming CD4^+^ T_FH_ cells, our data suggest that the migratory cDC play a more dominant role after 5-A-RU/MG administration, indicated by the preferential upregulation of co-stimulatory receptors on this population. Thus, the interactions driving DC activation most likely took place in the lung where local MAIT cells are present and capable of interacting with 5-OP-RU-loaded DC patrolling the lung tissue.

Earlier in vitro studies had indicated that MAIT cells can support B cell responses through production of interleukin (IL)-21, leading to preferential production of IgG^17,47^. Furthermore, cultures containing MAIT cells were also shown to exhibit enhanced memory B cell differentiation^48^. In a mouse model of lupus, MAIT cells were shown to directly enhance autoantibody production, GC formation, and T_FH_ cell responses in an MR1-dependent manner^47^. While other innate-like T cells such as NKT cells can adopt a T_FH_ phenotype and provide cognate help to B cells^25,26,49–51^, we were unable to identify a MAIT_FH_ population in the mLNs after using 5-A-RU/MG as an adjuvant in our study. In addition, MAIT cells were unable to support B cell immunity in B6Aa0^−/−^ mice which lack conventional CD4^+^ T_FH_ cells. This contrasts with studies using NKT agonists, where mice lacking conventional CD4^+^ T_FH_ cells generate a comparable antibody responses to WT mice^52^. However, a population of MAIT_FH_ (CXCR5^+^) has been identified in vivo following i.n. *Salmonella enterica* Serovar Typhimurium challenge, but not after 5-A-RU/MG stimulation, which was in line with our observations^53^. A role for such MAIT_FH_ in mice requires further investigation. While Bcl6^fl/fl^Cre^CD4^ mice that are deficient in CD4^+^ T_FH_ cells have been widely used to assess the role and contribution of conventional T_FH_, these mice exhibit a deficiency in MAIT cells, which require Bcl6 for their development^54^. Nevertheless, our data suggest that MAIT cells contribute to the humoral response through an indirect non-cognate role, involving activation of DC to prime conventional CD4^+^ T_FH_ cells. A similar non-cognate role has been described for NKT cells^55^ (in addition to generation of NKT_FH_).

While our study has focused on the “cellular adjuvant” properties of MAIT cells, it has previously been reported that MAIT cells can be harnessed as useful immune effectors in their own right. Prophylactic mucosal delivery of 5-OP-RU in mice has been shown to induce expansion of MAIT cell populations in the lung, leading to enhanced protection against subsequent pulmonary infection with *Legionella*^8^. Treatment with 5-OP-RU can also enhance control and clearance of *Mycobacterium tuberculosis* in mice^56^. However, in both studies, co-administration with a TLR agonist was critical for MAIT cells to differentiate into effector cells capable of producing IL-17 and IFNγ^8,56^. Activation of MAIT cells can also contribute to antiviral responses^21–23^, which we observed in mice pre-treated with 5-A-RU/MG alone before IAV challenge. However, by including virus antigen in the prophylactic treatment, the additional cellular adjuvant activities of MAIT cells were exploited, provoking an effective adaptive immune response that helped to hasten virus clearance, as indicated by improved weight gain. It remains to be established how long the two activities are maintained after administration, and whether additional TLR ligation will improve the response.

As the MAIT cell/MR1 axis is also observed in humans, MR1-binding agonists potentially represent a new class of adjuvants to be used in vaccines to initiate humoral immunity in the respiratory mucosa. However, recent studies have suggested that MAIT cell phenotype and function can differ between species, which may need to be considered when extending our observations to the clinic. For example, while treatment of chronically infected mice with 5-OP-RU (and TLR ligands) enhanced *Mycobacterium tuberculosis* clearance^56^, this was not observed in non-human primates (NHP); in fact, treatment rendered these animals more susceptible to infection^57^. Probing into the mechanism for this discrepancy revealed that, over time, MAIT cells in NHP expressed an exhausted phenotype, with high levels of inhibitory receptor PD-1 expression and reduced levels of IFNγ, IL-17 and granulocyte-macrophage colony-stimulating factor (GM-CSF)^57^. Similarly, transcriptional analysis of MAIT cells isolated from human BAL samples showed that this population is composed of heterogenous populations that include a subset of anti-inflammatory cells expressing a transcriptional profile associated with wound healing and genes encoding inhibitory receptors (i.e. T-cell immunoglobulin and mucin domain-containing protein-3 (TIM-3))^14^. Further, in vitro studies show triggering of MAIT TCRs preferentially induce tissue-repair programs^58^. Whether MAIT cells can promote T_FH_ differentiation and humoral immunity through vaccination in humans is therefore not guaranteed. However, in vitro studies showing that activated human MAIT cells can activate DCs is promising in this regard^15^.

In summary, this work identifies a role for MAIT cells as a cellular adjuvant to promote activation of local DCs, which ultimately enhance humoral responses in the respiratory tract. The data presented here provide good rationale to explore whether MAIT cells or the factors provided by MAIT cells can be used in a clinical setting to aid in the development of mucosal vaccines.

## Methods

### Mice

Mice were bred and housed at the Victoria University of Wellington Small Animal Facility and Malaghan Institute of Medical Research Biomedical Research Unit (Wellington, New Zealand). C57BL/6 (designated WT) mice were originally obtained from The Jackson Laboratory (Bar Harbor, ME). MR1^−/−^ mice that lack the α1 and α2 domains of the gene encoding MR1, were kindly provided by the late Vincenzo Cerundolo (Weatherall Institute, Oxford, UK). Nur77-GFP transgenic mice exhibit GFP expression under control of *Nr4al* region within the BAC transgene, consistent with endogenous Nur77 and antigen-receptor stimulation. Nur77-GFP mice were obtained from The Jackson Laboratory (Bar Harbor, ME). B6Aa0^−/−^ mice were created on a C57BL/6 background by targeting a mutation to the Aa gene encoding MHCII, provided by the Biological Research Laboratories Ltd (Wolferstrasse, Switzerland). Mice aged 8 to 12 weeks were used in experiments, after approval by the Victoria University of Wellington Animal Ethics Committee.

### Synthesis of 5-amino-6-D-ribitylaminouracil (5-A-RU)

Sodium dithionite (nominally 86% by redox titration, 84.0 mg, nominally 6.0 equiv) was added to a suspension of 5-nitroso-6-(D-ribitylamino)uracil (20.4 mg, 0.069 mmol) in sterile water (1.5 mL, Sigma-Aldrich). The mixture was briefly sonicated until the nitrosouracil dissolved and a colourless solution was obtained^59^.

### Immunisations

Intranasal procedures were performed in 30 μL volumes on ketamine/xylazine sedated mice. For single-dose assays, mice received either PBS or 5-A-RU/MG (MG; Sigma-Aldrich). For OVA-specific prime-boost assays, mice received either 5 nmol Endo Grade® Ovalbumin protein (Sigma-Aldrich) alone, or mixed with 5-A-RU/MG (75 nmol 5-A-RU mixed with 750 nmol MG). Three homologous immunisations were performed at two-week intervals. For HA-specific prime-boost assays, mice received either PBS, 5-A-RU/MG alone, 10 μg of IAV HA (A/X-31) protein alone, or IAV HA protein mixed with 5-A-RU/MG. As a benchmark, 10 μg of IAV HA was mixed with AddaVax (InvivoGen) at 1:1 v/v and delivered subcutaneously into the right and left flanks in 50 μL volumes according to the same prime-boost regimen as above.

### Tissue preparation and flow cytometry

For DC preparation, mLNs were collected and digested with DNase I and Liberase TL (Sigma-Aldrich) for 25 min at 37 °C. For lung preparation, lungs were perfused prior to collection and finely chopped before digestion with DNase I and Liberase TL for 45 min at 37 °C. Cells were passed through a 70 μm cell strainer and washed in preparation for staining. Single cell suspensions were stained for viability with Zombie NIR/Aqua™ (BioLegend) or BD Horizon™ Fixable Viability Stain 700 (BD Biosciences), before incubation with anti-mouse CD16/32 (clone 2.4G2), to block any non-specific antibody binding. For labelling OVA-specific T_FH_ cells, prior to surface staining, cells were incubated with two MHCII-OVA tetramers (containing I-A(b)HAAHAEINEA and I-A(b)AAHAEINEA epitopes both labelled with PE; NIH Tetramer Core Facility) for 30 min at RT. For labelling OVA-specific B cells, cells were incubated with biotinylated OVA protein (made in-house) for 30 min at 4 °C. Cells were stained in a separate step for chemokine receptor CXCR5 (2G8) at 37 °C for 30 min. For staining of surface molecules, cells were incubated with cocktails of fluorescent antibodies specific for: CD11c (N418, HL3), MHCII (I-A/I-E M5/114.15.2, 3JP), CD86 (GL-1), CD80 (16-10A1), CD64 (X54-5/7.1, X54-5/7.1), CD11b (M1/70), SIRP1α (P84), PDL-1 (MIH5), CD69 (H1.2F3), B220 (RA306B2, RA3-6B2), TCRβ (H57-597), CD4 (GK1.5), CD44 (IM7), ICOS (15F9), FR4 (12A5), CD25 (PC61), CD40 (3/23), PD-1 (29F.1A12), GL7 (GL7), IgD (11-26c.2a), CD40L (MR1), CD45.2 (A20), and MR1-5OPRU tetramer (NIH Tetramer Core). For intranuclear transcription factor staining, cells were fixed and permeabilized using the FOXP3 Transcription Factor Staining Buffer Set (Life Technologies), before staining with Bcl6 (K112-91), or Ki67 (11F6). Data were acquired on a LSRFortessa Flow Cytometer (BD) or Cytek™ Aurora (Cytek Biosciences) and analysed using FlowJo version 9.6.2 (Tree Star).

### Detection of serum IgG by ELISA

For detection of antigen-specific serum IgG, microtitre plates (Maxisorp, Nunc) were incubated overnight with 20 μg/mL Endo Grade® Ovalbumin (Sigma-Aldrich), or 10 μg/mL IAV HA in bicarbonate buffer. 10% FBS was applied to prevent non-specific binding, prior to application of serially diluted serum samples. A final incubation was performed with a variety of Ig isotypes; IgG_TOTAL_ (G21040), IgG1 (A10551) IgG2a/c (PA129288), IgG2b (M32407), IgG3 (M32707) conjugated to HRP (Life Technologies). Tetramethylbenzidine (TMB; BD Biosciences) and 2M H_2_SO_4_ (Sigma-Aldrich) was applied to develop and stop reactions, before immediately being read at 450 nm. Wash steps were performed 5x between each incubation with 10% PBS-Tween (Sigma-Aldrich).

### Detection of serum IgE by ELISA

Serum IgE antibodies were measured using a mouse IgE ELISA kit (BD Biosciences) according to manufacturer’s instructions.

### Detection of BALF IgA by ELISA

BALF was collected by inserting a catheter into the trachea. 0.6 mL of 10% FCS PBS was injected and aspirated three times. Approximately 0.5 mL of BALF was recovered from each mouse and supernatant was collected following centrifugation. Aliquots of BALF were stored under −80 °C. IgA antibodies in the BALF were measured using a mouse Total IgA ELISA kit (Thermo Fisher Scientific) according to manufacturer’s instructions with the exception of the coating antigen. For detection of antigen-specific IgA, microtitre plates (Maxisorp, Nunc) were incubated overnight with either 20 μg/mL Endo Grade® Ovalbumin (Sigma-Aldrich), or 10 μg/mL IAV HA in bicarbonate buffer.

### Preparation of haemagglutinin (HA) protein

HEK293S GnTI-cells were transfected using polyethylenimine with the cDNA encoding for influenza A (A/X-31) HA ectodomain fused to the Fc region of human IgG as well as an empty vector plasmid (pcDNA3.1) conferring G418 resistance. After 48 hours, cells were selected in DMEM supplemented with 5% FBS and 500 μg/mL G418. Resistant clones were isolated using Pyrex cloning rings (Corning) and levels of expression of the Fc-HA protein were tested via western blot. Best expressing clones were used for large-scale protein production, which was performed in cell culture flasks (Nunc™ TripleFlask^TM)^ with regular collection and replenishment of cell culture medium, containing 2-5% FBS. Secreted proteins were purified by affinity chromatography, using CaptivA™ PriMAB rProtein A Affinity Resin (Repligen). Resin was washed (50 mM Tris pH 7.4, 450 mM NaCl), equilibrated (50 mM Tris pH 7.4, 150 mM NaCl, 1mM DTT) and the HA portion was cleaved by HRV-3C protease (50 mM Tris pH 7.4, 150 mM NaCl, 1mM DTT, 10μg/mL HRV-3C protease) overnight. Purified protein was buffer exchanged into PBS and concentrated using a Vivaspin concentrator (Sartorius-Stedim) to 3-5 mg/mL and flash-frozen with liquid nitrogen and stored at −80 °C.

### Influenza challenge

Three weeks after the final third boost immunisation, ketamine/xylazine sedated mice were challenged with a 50 μL injection containing LD50 2HAU A/X-31 (A/HK/8/68;ATCC). Mice were weighed daily for two weeks post-infection. Mice that lost more than 70% of initial body weight were euthanised.

### αCD40L monoclonal antibody blockade

Mice were administered 250-500 μg of anti-mouse CD40L (CD154) (clone MR-1; BioXCell) intraperitoneally as indicated in the relevant figure legends. Control mice received 250-500 μg of Armenian Hamster IgG (BioXCell).

### Statistical analyses

Statistical tests were performed using the GraphPad Prism software (Version 8, GraphPad, CS, USA). Data are represented as mean ± SD. Statistical analysis of p<0.05 was considered significant, determined by unpaired two-tailed Student’s t-test, log-rank (Mantel-cox) test, one and two-way analysis of variance (ANOVA) with Tukey’s multiple-comparison test, as specified in the relevant figure legends.

### Data availability

The authors declare that the data supporting the findings of this study are available within the paper and its supplementary information files are available from the authors upon reasonable requests.

## Acknowledgments

We thank the Malaghan Institute of Medical Research Biomedical Research Unit and the Victoria University of Wellington Small Animal Facility for provision and husbandry of mice. We thank the Malaghan Institute of Medical Research Hugh Green Cytometry Core for maintenance of flow cytometers. We thank the NIH Tetramer Core Facility (contract number 75N93020D00005) for providing MR1-5-OP-RU and MHCII-OVA tetramers. We thank Prof. Franca Ronchese for the expert opinion provided on this study, as well as Dr. Olivia Burn, Dr. Alissa Cait and Dr. David O’Sullivan for reviewing our manuscript. This research was funded by a Health Research Council of New Zealand Project Grant to L.M.C, Malaghan Institute of Medical Research New Zealand and Research for Life New Zealand. T.E.P. was supported by a Victoria University of Wellington Doctoral Research Scholarship. K.H.B. was supported by a Victoria University of Wellington Masters by Thesis Scholarship. I.F.H. was supported by the Thompson Family Foundation and Hugh Dudley Morgans Fellowship. L.M.C. was supported by a Sir Charles Hercus Health Research Fellowship from the Health Research Council of New Zealand.

## Author contributions

T.E.P, K.H.B, J.L.L, K.R.B, K.J.F, O.R.P, I.F.N.S and N.C.M performed experiments. A.M, T.B, B.C, D.C, M.S, G.F.P, I.F.H and L.M.C provided vital materials and expertise. T.E.P, K.H.B, J.L.L, G.F.P, I.F.H and L.M.C designed and analysed experiments, coordinated the study, and wrote the paper.

## Materials & Correspondence

Correspondence and requests for materials should be addressed to L.M.C.

## References

1. Li, M., Wang, Y., Sun, Y., Cui, H., Zhu, S. J. & Qiu, H. J. Mucosal vaccines: Strategies and challenges. Immunol. Lett. 217, 116–125 (2020).

2. Corthésy, B. Multi-faceted functions of secretory IgA at mucosal surfaces. Front. Immunol. 4, 185 (2013).

3. Carter, N. J. & Curran, M. P. Live attenuated influenza vaccine (FluMist®; FluenzTM): a review of its use in the prevention of seasonal influenza in children and adults. Drugs. 71, 1591–1622 (2011).

4. Lindsey, B. B., Jagne, Y. J., Armitage, E. P., Singanayagam, A., Sallah, H. J., Drammeh, S., Senghore, E., Mohammed, N. I., Jeffries, D., Höschler, K., Tregoning, J. S., Meijer, A., Clarke, E., Dong, T., Barclay, W., Kampmann, B. & de Silva, T.I. Effect of a Russian-backbone live-attenuated influenza vaccine with an updated pandemic H1N1 strain on shedding and immunogenicity among children in The Gambia: an open-label, observational, phase 4 study. Lancet Respir. Med. 7, 665–676 (2019).

5. Calzas, C. & Chevalier, C. Innovative mucosal vaccine formulations against influenza a virus infections. Front. Immunol. 10, 1605 (2019).

6. Treiner, E., Duban, L., Bahram, S., Radosavljevic, M., Wanner, V., Tilloy, F., Affaticati, P., Gilfillan, S. & Lantz, O. Selection of evolutionarily conserved mucosal-associated invariant T cells by MR1. Nature. 422, 164–169 (2003).

7. Rahimpour, A., Koay, H. F., Enders, A., Clanchy, R., Eckle, S. B. G., Meehan, B., Chen, Z., Whittle, B., Liu, L., Fairlie, D. P., Goodnow, C. C., McCluskey, J., Rossjohn, J., Uldrich, A. P., Pellicci, D. G. & Godfrey, D. I. Identification of phenotypically and functionally heterogeneous mouse mucosal-associated invariant T cells using MR1 tetramers. J. Exp. Med. 212, 1095–1108 (2015).

8. Wang, H., D’Souza, C., Lim, X. Y., Kostenko, L., Pediongco, T. J., Eckle, S. B. G., Meehan, B. S., Shi, M., Wang, N., Li, S., Liu, L., Mak, J. Y. W., Fairlie, D. P., Iwakura, Y., Gunnersen, J. M., Stent, A. W., Godfrey, D. I., Rossjohn, J., Westall, G. P., et al. MAIT cells protect against pulmonary Legionella longbeachae infection. Nat. Commun. 9, 1–15 (2018).

9. Corbett, A. J., Eckle, S. B. G., Birkinshaw, R. W., Liu, L., Patel, O., Mahony, J., Chen, Z., Reantragoon, R., Meehan, B., Cao, H., Williamson, N. A., Strugnell, R. A., Van Sinderen, D., Mak, J. Y. W., Fairlie, D. P., Kjer-Nielsen, L., Rossjohn, J. & McCluskey, J. T-cell activation by transitory neo-antigens derived from distinct microbial pathways. Nature. 509, 361–365 (2014).

10. Kjer-Nielsen, L., Patel, O., Corbett, A. J., Le Nours, J., Meehan, B., Liu, L., Bhati, M., Chen, Z., Kostenko, L., Reantragoon, R., Williamson, N. A., Purcell, A. W., Dudek, N. L., McConville, M. J., O’Hair, R. A. J., Khairallah, G. N., Godfrey, D. I., Fairlie, D. P., Rossjohn, J., et al. MR1 presents microbial vitamin B metabolites to MAIT cells. Nature. 491, 717–723 (2012).

11. Patel, O., Kjer-Nielsen, L., Le Nours, J., Eckle, S. B. G., Birkinshaw, R., Beddoe, T., Corbett, A. J., Liu, L., Miles, J. J., Meehan, B., Reantragoon, R., Sandoval-Romero, M. L., Sullivan, L. C., Brooks, A. G., Chen, Z., Fairlie, D. P., McCluskey, J. & Rossjohn, J. Recognition of vitamin B metabolites by mucosal-associated invariant T cells. Nat. Commun. 4, 1–9 (2013).

12. Meierovics, A., Yankelevich, W. J. C. & Cowley, S. C. MAIT cells are critical for optimal mucosal immune responses during in vivo pulmonary bacterial infection. Proc. Natl. Acad. Sci. U. S. A. 110, E3119–E3128 (2013).

13. Chen, Z., Wang, H., Souza, D. ‘, Sun, S., Kostenko, L., Eckle, S. B. G., Meehan, B. S., Jackson, D. C., Strugnell, R. A., Cao, H., Wang, N., Fairlie, D. P., Liu, L., Godfrey, D. I., Rossjohn, J., McCluskey, J., Corbett, A. J., D’Souza, C., Sun, S., et al. Mucosal-associated invariant T-cell activation and accumulation after in vivo infection depends on microbial riboflavin synthesis and co-stimulatory signals. Mucosal. Immunol. 10, 58–68 (2017).

14. Hinks, T. S. C., Marchi, E., Jabeen, M., Olshansky, M., Kurioka, A., Pediongco, T. J., Meehan, B. S., Kostenko, L., Turner, S. J., Corbett, A. J., Chen, Z., Klenerman, P. & McCluskey, J. Activation and In Vivo Evolution of the MAIT Cell Transcriptome in Mice and Humans Reveals Tissue Repair Functionality. Cell Rep. 28, 3249–3262 (2019).

15. Salio, M., Gasser, O., Gonzalez-lopez, C., Martens, A., Veerapen, N., Gileadi, U., Jacob, G., Napolitani, G., Anderson, R., Painter, G., Besra, G. S., Hermans, I. F. & Cerundolo, V. Activation of Human Mucosal-Associated Invariant T Cells Induces CD40L-Dependent Maturation of Monocyte-Derived and Primary Dendritic Cells. J. Immunol. 199, 2631–2638 (2017).

16. Crotty, S. T Follicular Helper Cell Biology: A Decade of Discovery and Diseases. mmunity. 50, 1132–1148 (2019).

17. Bennett, M. S., Trivedi, S., Iyer, A. S., Hale, J. S. & Leung, D. T. Human mucosal-associated invariant T (MAIT) cells possess capacity for B cell help. J. Leukoc. Biol. 102, 1261–1269 (2017).

18. Macpherson, A. J., McCoy, K. D., Johansen, F.-E. & Brandtzaeg, P. The immune geography of IgA induction and function. Mucosal. Immunol. 1, 11–22 (2008).

19. Russell, R. J., Kerry, P. S., Stevens, D. J., Steinhauer, D. A., Martin, S. R., Gamblin, S. J. & Skehel, J. J. Structure of influenza hemagglutinin in complex with an inhibitor of membrane fusion. Proc. Natl. Acad. Sci. U. S. A. 105, 17736 (2008).

20. Onodera, T., Takahashi, Y., Yokoi, Y., Ato, M., Kodama, Y., Hachimura, S., Kurosaki, T. & Kobayashi, K. Memory B cells in the lung participate in protective humoral immune responses to pulmonary influenza virus reinfection. Proc. Natl. Acad. Sci. U. S. A. 109, 2485–2490 (2012).

21. Wilgenburg, B. van, Loh, L., Chen, Z., Pediongco, T. J., Wang, H., Shi, M., Zhao, Z., Koutsakos, M., Nüssing, S., Sant, S., Wang, Z., D’Souza, C., Jia, X., Almeida, C. F., Kostenko, L., Eckle, S. B. G., Meehan, B. S., Kallies, A., Godfrey, D. I., et al. MAIT cells contribute to protection against lethal influenza infection in vivo. Nat. Commun. 9, 1–9 (2018).

22. Van Wilgenburg, B., Scherwitzl, I., Hutchinson, E. C., Leng, T., Kurioka, A., Kulicke, C., De Lara, C., Cole, S., Vasanawathana, S., Limpitikul, W., Malasit, P., Young, D., Denney, L., Moore, M. D., Fabris, P., Giordani, M. T., Oo, Y. H., Laidlaw, S. M., Dustin, L. B., et al. MAIT cells are activated during human viral infections. Nat. Commun. 7, 1– 11 (2016).

23. Ussher, J. E., Bilton, M., Attwod, E., Shadwell, J., Richardson, R., de Lara, C., Mettke, E., Kurioka, A., Hansen, T. H., Klenerman, P. & Willberg, C. B. CD161++ CD8+ T cells, including the MAIT cell subset, are specifically activated by IL-12+IL-18 in a TCR-independent manner. Eur. J. Immunol. 44, 195–203 (2014).

24. Moran, A. E., Holzapfel, K. L., Xing, Y., Cunningham, N. R., Maltzman, J. S., Punt, J. & Hogquist, K. A. T cell receptor signal strength in Treg and iNKT cell development demonstrated by a novel fluorescent reporter mouse. J. Exp. Med. 208, 1279–1289 (2011).

25. Chang, P. P., Barral, P., Fitch, J., Pratama, A., Ma, C. S., Kallies, A., Hogan, J. J., Cerundolo, V., Tangye, S. G., Bittman, R., Nutt, S. L., Brink, R., Godfrey, D. I., Batista, F. D. & Vinuesa, C. G. Identification of Bcl-6-dependent follicular helper NKT cells that provide cognate help for B cell responses. Nat. Immunol. 13, 35–43 (2012).

26. King, I. L., Fortier, A., Tighe, M., Dibble, J., Watts, G. F. M., Veerapen, N., Haberman, A. M., Besra, G. S., Mohrs, M., Brenner, M. B. & Leadbetter, E. A. Invariant natural killer T cells direct B cell responses to cognate lipid antigen in an IL-21-dependent manner. Nat. Immunol. 13, 44–50 (2012).

27. Pennock, N. D., White, J. T., Cross, E. W., Cheney, E. E., Tamburini, B. A. & Kedl, R. M. T cell responses: naïve to memory and everything in between. Adv. Physiol. Educ. 37, 273 (2013).

28. Krishnaswamy, J. K., Gowthaman, U., Zhang, B., Mattsson, J., Szeponik, L., Liu, D., Wu, R., White, T., Calabro, S., Xu, L., Collet, M. A., Yurieva, M., Alsén, S., Fogelstrand, P., Walter, A., Heath, W. R., Mueller, S. N., Yrlid, U., Williams, A., et al. Migratory CD11b + conventional dendritic cells induce T follicular helper cell-dependent antibody responses. Sci. Immunol. 2, (2017).

29. Krishnaswamy, J. K., Alsén, S., Yrlid, U., Eisenbarth, S. C. & Williams, A. Determination of T Follicular Helper Cell Fate by Dendritic Cells. Frontiers in immunology. 9, 2169 (2018).

30. Bouteau, A., Kervevan, J., Su, Q., Zurawski, S. M., Contreras, V., Dereuddre-Bosquet, N., Le Grand, R., Zurawski, G., Cardinaud, S., Levy, Y. & Igyártó, B. Z. DC subsets regulate humoral immune responses by supporting the differentiation of distinct TFH cells. Front. Immunol. 10, 1134 (2019).

31. Pooley, J. L., Heath, W. R. & Shortman, K. Cutting Edge: Intravenous Soluble Antigen Is Presented to CD4 T Cells by CD8-Dendritic Cells, but Cross-Presented to CD8 T Cells by CD8+ Dendritic Cells. J. Immunol. 166, 5327–5330 (2001).

32. Hildner, K., Edelson, B. T., Purtha, W. E., Diamond, M., Matsushita, H., Kohyama, M., Calderon, B., Schraml, B. U., Unanue, E. R., Diamond, M. S., Schreiber, R. D., Murphy, T. L. & Murphy, K. M. Batf3 Deficiency Reveals a Critical Role for CD8α+ Dendritic Cells in Cytotoxic T Cell Immunity. Science. 322, 1097 (2008).

33. Durai, V. & Murphy, K. M. Functions of Murine Dendritic Cells. Immunity. 45, 719–736 (2016).

34. Cella, M., Scheidegger, D., Palmer-Lehmann, K., Lane, P., Lanzavecchia, A. & Alber, G. Ligation of CD40 on dendritic cells triggers production of high levels of interleukin-12 and enhances T cell stimulatory capacity: T-T help via APC activation. J. Exp. Med. 184, 747–752 (1996).

35. Lanzavecchia, A. Immunology. Licence to kill. Nature. 393, 413–414 (1998).

36. Bennett, S. R. M., Carbone, F. R., Karamalis, F., Miller, J.F.A.P. & Heath, W. R. Induction of a CD8+ Cytotoxic T Lymphocyte Response by Cross-priming Requires Cognate CD4+ T Cell Help. J. Exp. Med. 186, 65 (1997).

37. Ridge, J. P., Di Rosa, F. & Matzinger, P. A conditioned dendritic cell can be a temporal bridge between a CD4+ T-helper and a T-killer cell. Nature. 393, 474–478 (1998).

38. Schoenberger, S. P., Toes, R. E. M., Van Dervoort, E. I. H., Offringa, R. & Melief, C. J. M. T-cell help for cytotoxic T lymphocytes is mediated by CD40–CD40L interactions. Nature. 393, 480–483 (1998).

39. Nishimura, T., Kitamura, H., Iwakabe, K., Yahata, T., Ohta, A., Sato, M., Takeda, K., Okumura, K., Van Kaer, L., Kawano, T., Taniguchi, M., Nakui, M., Sekimoto, M. & Koda, T. The interface between innate and acquired immunity: glycolipid antigen presentation by CD1d-expressing dendritic cells to NKT cells induces the differentiation of antigen-specific cytotoxic T lymphocytes. Int. Immunol. 12, 987–994 (2000).

40. Hermans, I. F., Silk, J. D., Gileadi, U., Salio, M., Mathew, B., Ritter, G., Schmidt, R., Harris, A. L., Old, L. & Cerundolo, V. NKT Cells Enhance CD4 + and CD8 + T Cell Responses to Soluble Antigen In Vivo through Direct Interaction with Dendritic Cells. J. Immunol. 171, 5140–5147 (2003).

41. Inoue, S. I., Niikura, M., Takeo, S., Mineo, S., Kawakami, Y., Uchida, A., Kamiya, S. & Kobayashi, F. Enhancement of dendritic cell activation via CD40 ligand-expressing γδ T cells is responsible for protective immunity to Plasmodium parasites. Proc. Natl. Acad. Sci. U. S. A. 109, 12129–12134 (2012).

42. Kerkmann, M., Rothenfusser, S., Hornung, V., Towarowski, A., Wagner, M., Sarris, A., Giese, T., Endres, S. & Hartmann, G. Activation with CpG-A and CpG-B Oligonucleotides Reveals Two Distinct Regulatory Pathways of Type I IFN Synthesis in Human Plasmacytoid Dendritic Cells. J. Immunol. 170, 4465–4474 (2003).

43. Lind, E. F., Ahonen, C. L., Wasiuk, A., Kosaka, Y., Becher, B., Bennett, K. A. & Noelle, R. J. Dendritic cells require the NF-kappaB2 pathway for cross-presentation of soluble antigens. J. Immunol. 181, 354–363 (2008).

44. Kobayashi, T., Walsh, P. T., Walsh, M. C., Speirs, K. M., Chiffoleau, E., King, C. G., Hancock, W. W., Caamano, J. H., Hunter, C. A., Scott, P., Turka, L. A. & Choi, Y. TRAF6 is a critical factor for dendritic cell maturation and development. Immunity. 19, 353–363 (2003).

45. Kawabe, T., Naka, T., Yoshida, K., Tanaka, T., Fujiwara, H., Suematsu, S., Yoshida, N., Kishimoto, T. & Kikutani, H. The immune responses in CD40-deficient mice: impaired immunoglobulin class switching and germinal center formation. Immunity. 1, 167–178 (1994).

46. Hilligan, K. L. & Ronchese, F. Antigen presentation by dendritic cells and their instruction of CD4+ T helper cell responses. Cell. Mol. Immunol. 17, 587–599 (2020).

47. Murayama, G., Chiba, A., Suzuki, H., Nomura, A., Mizuno, T., Kuga, T., Nakamura, S., Amano, H., Hirose, S., Yamaji, K., Suzuki, Y., Tamura, N. & Miyake, S. A Critical Role for Mucosal-Associated Invariant T Cells as Regulators and Therapeutic Targets in Systemic Lupus Erythematosus. Front. Immunol. 10, 2681 (2019).

48. Rahman, M. A., Ko, E. J., Bhuyan, F., Enyindah-Asonye, G., Hunegnaw, R., Helmold Hait, S., Hogge, C. J., Venzon, D. J., Hoang, T. & Robert-Guroff, M. Mucosal-associated invariant T (MAIT) cells provide B-cell help in vaccinated and subsequently SIV-infected Rhesus Macaques. Sci. Rep. 10, 10060 (2020).

49. Galli, G., Pittoni, P., Tonti, E., Malzone, C., Uematsu, Y., Tortoli, M., Maione, D., Volpini, G., Finco, O., Nuti, S., Tavarini, S., Dellabona, P., Rappuoli, R., Casorati, G. & Abrignani, S. Invariant NKT cells sustain specific B cell responses and memory. Proc. Natl. Acad. Sci. U. S. A. 104, 3984–3989 (2007).

50. Leadbetter, E. A., Brigl, M., Illarionov, P., Cohen, N., Luteran, M. C., Pillai, S., Besra, G. S. & Brenner, M. B. NK T cells provide lipid antigen-specific cognate help for B cells. Proc. Natl. Acad. Sci. U. S. A. 105, 8339–8344 (2008).

51. Barral, P., Eckl-Dorna, J., Harwood, N. E., De Santo, C., Salio, M., Illarionov, P., Besra, G. S., Cerundolo, V. & Batista, F. D. B cell receptor-mediated uptake of CD1d-restricted antigen augments antibody responses by recruiting invariant NKT cell help in vivo. Proc. Natl. Acad. Sci. U. S. A. 105, 8345–8350 (2008).

52. Tonti, E., Fedeli, M., Napolitano, A., Iannacone, M., von Andrian, U. H., Guidotti, L. G., Abrignani, S., Casorati, G. & Dellabona, P. Follicular Helper NKT Cells Induce Limited B Cell Responses and Germinal Center Formation in the Absence of CD4 + T Cell Help. J. Immunol. 188, 3217–3222 (2012).

53. Jensen, O., Trivedi, S., Meier, J. D., Fairfax, K. C., Hale, J. S. & Leung, D. T. A subset of follicular helper-like MAIT cells can provide B cell help and support antibody production in the mucosa. Sci. Immunol. 7, (2022).

54. Gioulbasani, M., Galaras, A., Grammenoudi, S., Moulos, P., Dent, A. L., Sigvardsson, M., Hatzis, P., Kee, B. L. & Verykokakis, M. The transcription factor BCL-6 controls early development of innate-like T cells. Nat. Immunol. 21, 1058–1069 (2020).

55. Tonti, E., Galli, G., Malzone, C., Abrignani, S., Casorati, G. & Dellabona, P. NKT-cell help to B lymphocytes can occur independently of cognate interaction. Blood. 113, 370– 376 (2009).

56. Sakai, S., Kauffman, K. D., Oh, S., Nelson, C. E., Barry, C. E. & Barber, D. L. MAIT cell-directed therapy of Mycobacterium tuberculosis infection. Mucosal. Immunol. 14, 199–208 (2020).

57. Sakai, S., Lora, N. E., Kauffman, K. D., Dorosky, D. E., Oh, S., Namasivayam, S., Gomez, F., Fleegle, J. D., Bleach, J. L., Butler, A. L., Dayao, E. K., Piazza, M. K., Repoli, K. M., Slone, B. Y., Sutphin, M. K., Vatthauer, A. M., Walker, A. M., Weiner, D. M., Woodcock, M. J., et al. Functional inactivation of pulmonary MAIT cells following 5-OP-RU treatment of non-human primates. Mucosal. Immunol. 14, 1055–1066 (2021).

58. Leng, T., Akther, H. D., Hackstein, C. P., Powell, K., King, T., Friedrich, M., Christoforidou, Z., McCuaig, S., Neyazi, M., Arancibia-Cárcamo, C. V., Hagel, J., Powrie, F., Peres, R. S., Millar, V., Ebner, D., Lamichhane, R., Ussher, J., Hinks, T. S. C., Marchi, E., et al. TCR and Inflammatory Signals Tune Human MAIT Cells to Exert Specific Tissue Repair and Effector Functions. Cell. Rep. 28, 3077–3091 (2019).

59. Lange, J., Anderson, R. J., Marshall, A. J., Chan, S. T. S., Bilbrough, T. S., Gasser, O., Gonzalez-Lopez, C., Salio, M., Cerundolo, V., Hermans, I. F. & Painter, G. F. The Chemical Synthesis, Stability, and Activity of MAIT Cell Prodrug Agonists That Access MR1 in Recycling Endosomes. ACS Chem. Biol. 15, 437–445 (2020).

