## Supplementary materials for "MAIT cells activate dendritic cells to promote T follicular helper cell differentiation and humoral immunity"

**Supplementary Figures**

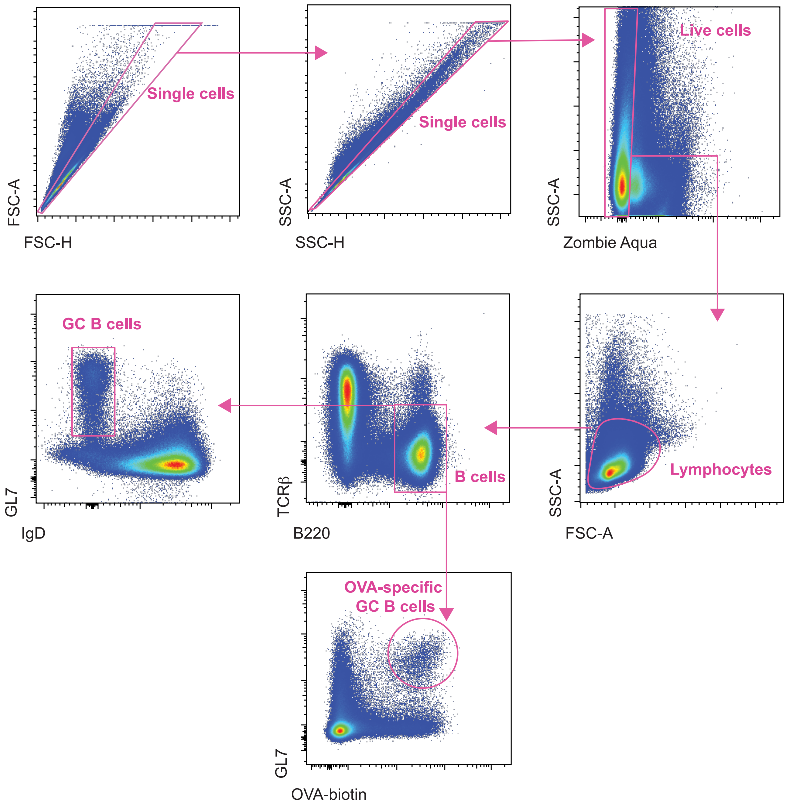


**Supplementary Figure 1. Flow cytometry gating strategy for GC B cells** Relates to Figure 1 and Figure 5. Murine GC B cells were identified by excluding doublets using forward scatter/side scatter properties; live lymphocytes were gated and further selected as $B220^{+}TCR\beta^{-}$ B cells; naïve B cells were excluded as $IgD^{-}$; $GL7^{+}$ cells were selected to denote GC activity and $OVA\text{-}biotin^{+}$ cells were gated for antigen-specificity.

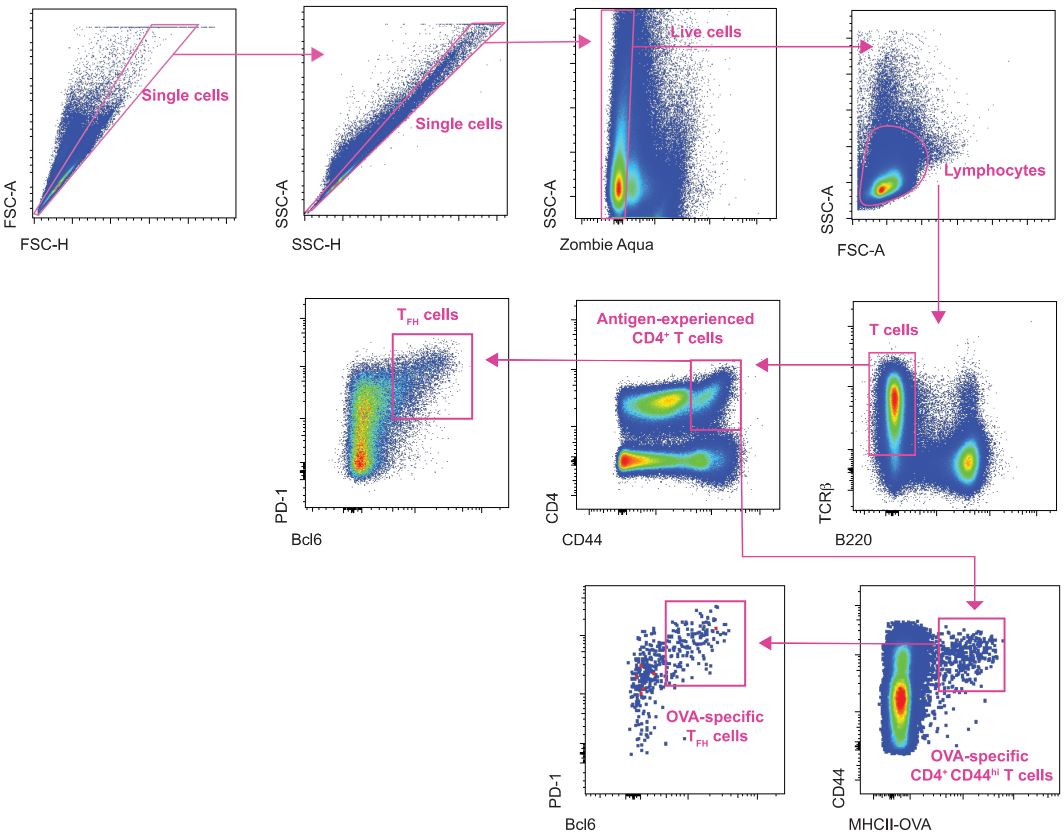


**Supplementary Figure 2. Flow cytometry gating strategy for $T_{FH}$ cells** Relates to Figure 4 and Figure 7. Murine $T_{FH}$ cells were identified by excluding doublets using forward scatter/side scatter properties; live lymphocytes were then gated and further selected as $TCR\beta^{+}B220^{-}$ T cells; antigen-experienced $CD4^{+}$ T cells were selected as $CD4^{+}CD44^{hi}$; TFH cells were either gated next as $Bcl6^{+}PD\text{-}1^{+}$, or first gated for antigen-specificity via two overlapping MHC class II OVA tetramers on the same fluorophore (MHCII-OVA) prior to gating for $Bcl6^{+}PD\text{-}1^{+}$ $T_{FH}$ markers.

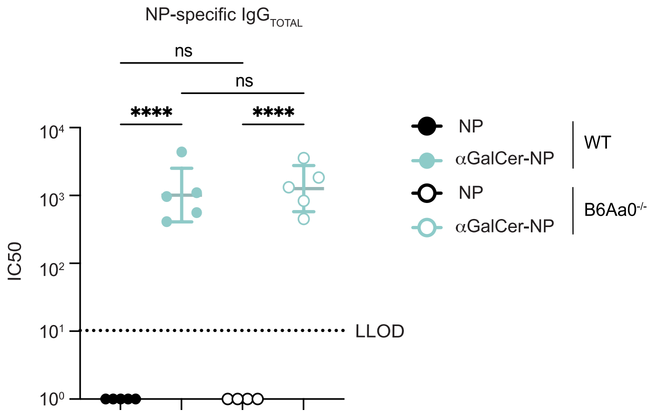


**Supplementary Figure 3. Antigen-specific antibody responses retained in B6Aa0$^{-/-}$ towards T-independent antigen** Relates to Figure 5. C57BL/6 **(**WT) or B6Aa0$^{-/-}$ mice were immunized intravenously (i.v.) with a single dose of either 2.5 nmol of hapten antigen 4-hydroxy-3-nitrophenyl (NP) alone, or in combination with 2.5 nmol of α-galactosylceramide (αGalCer). Six weeks later serum was collected for analysis. IC50 Log10 values from NP-specific serum IgG ELISA. LLOD was set at an IC50 of 10, IC50 values below zero were marked as an IC50 of one. Graphs depicted as mean ± SD. Data represent *n*=2 individual experiments, with *n*=4-5 mice per group. One-way ANOVA with Tukey's multiple comparisons test was performed with ns>0.05, ****$P$≤0.0001.

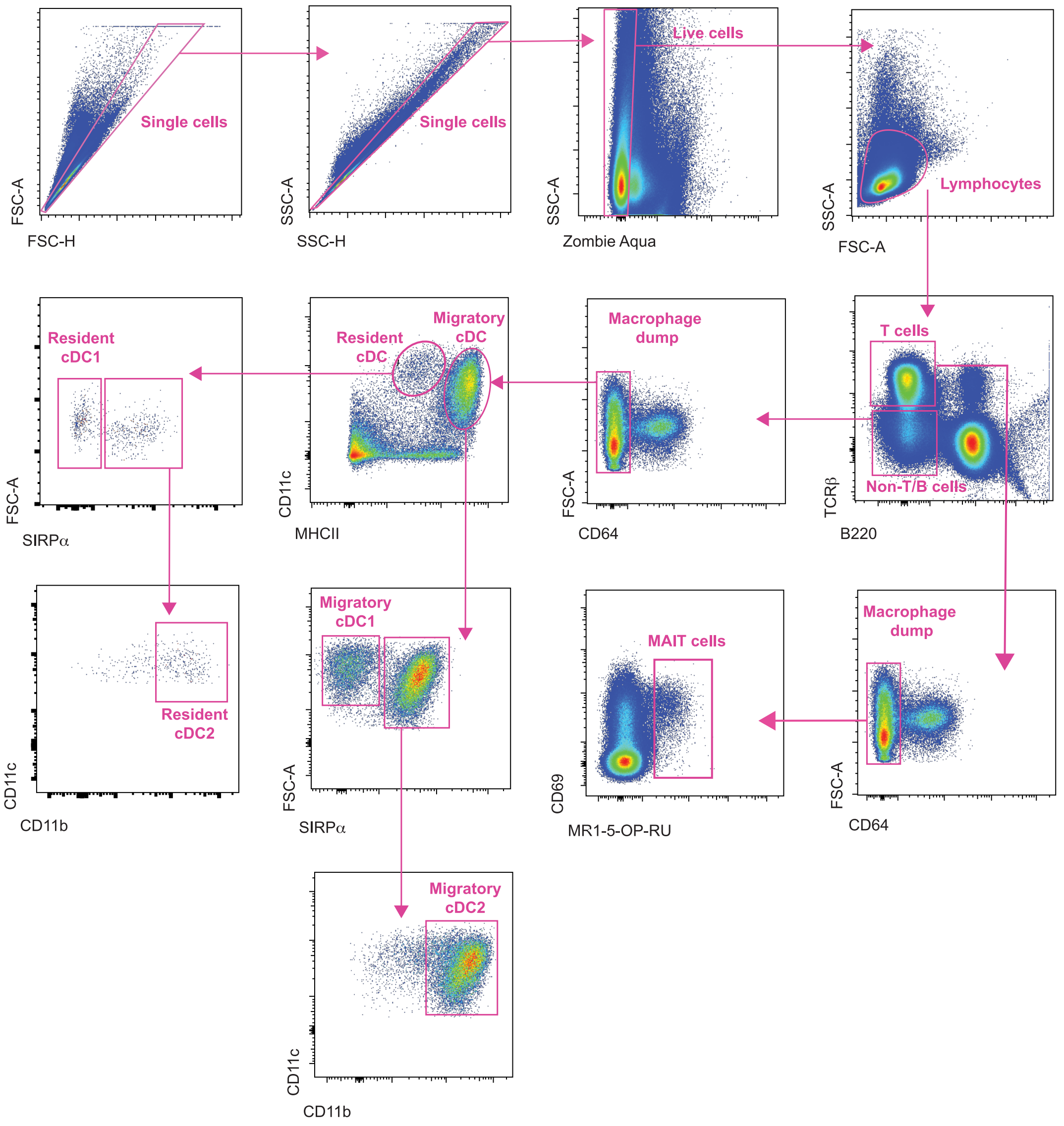


**Fig. S4. Flow cytometry gating for MAIT cells and DCs.** Relates to Figure 3 and Figure 6. Murine MAIT cells and DCs were identified by excluding doublets using forward scatter / side scatter properties; live lymphocytes were gated; TCRβ$^+$ T cells were selected; CD64$^+$ cells were excluded to remove macrophages and a fluorescent MR1-5-OP-RU tetramer was used to identify MAIT cells. To assess DCs, TCRβ$^+$B220$^+$ cells were excluded to remove T and B cells; CD64$^+$ cells were excluded to remove macrophages; CD11c$^+$MHCII$^{hi}$ cells were selected as migratory cDC, and CD11c$^+$MHCII$^{Int}$ cells were selected as resident cDC. Both migratory and resident cDC subsets could be further selected as CD11c$^+$SIRPα$^-$ cDC1s or CD11c$^+$CD11b$^+$SIRPα$^+$ cDC2s.

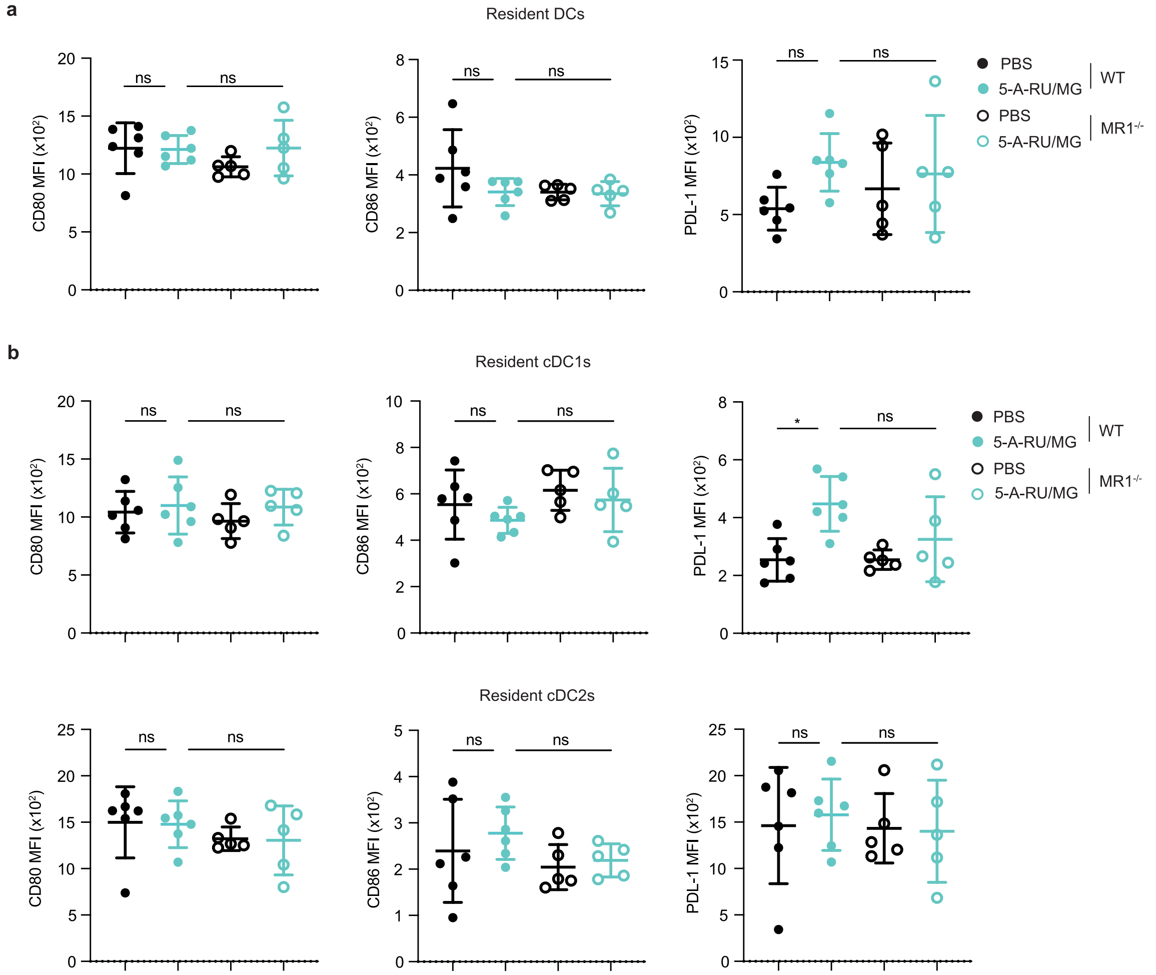


**Supplementary Figure 5. Resident cDC populations in the mLN are not activated following intranasal 5-A-RU/MG administration.** Relates to Figure 6. (**a**) WT or MR1$^{-/-}$ mice were administered i.n. with 75 nmol 5-A-RU and 750 nmol MG or PBS. 24 hours later mLN were harvested and stained for flow cytometry. (**a**) MFI of activation markers expressed on resident cDCs. (**b**) MFI of activation markers expressed on resident cDC1 and cDC2 populations. Graphs depicted as mean ± SD. Data representative of *n*=2 individual experiments, with *n*=5-6 mice per group. Statistical significance was determined by One-way ANOVA with Tukey's multiple comparisons test. ns>0.05, **P*≤0.05.

**Supplementary methods**

**αGalCer-NP synthesis**

Relates to Supplementary Figure 3. Briefly, 2-(4-((4-methoxybenzyl)oxy)-3-nitrophenyl)acetate was saponified with sodium hydroxide and the resulting acid was coupled with N-hydroxysuccinimide. This activated ester was then treated with methyl (((9H-fluoren-9-yl)methoxy)carbonyl)-L-lysinate followed by removal of the fluorenylmethyloxycarbonyl (Fmoc) group. The resulting lysine derivative was treated with tert-Boc-aminooxyacetic acid N-hydroxysuccinimide ester to afford the fully protected hapten. Trifluoroacetic acid-mediated removal of the tert-butyloxycarbonyl (Boc) and para-methoxybenzyl (PMB) groups gave the aminooxy-containing product which was conjugated to 6''-amino-6''-deoxy-alpha-galactosylceramide bearing the para-aminobenzyl-citrulline-valine-nonanone immolative linker to afford the adjuvanted hapten, αGalCer-NP.
